# Imaging the extracellular matrix in live tissues and organisms with a glycan-binding fluorophore

**DOI:** 10.1101/2024.05.09.593460

**Authors:** Antonio Fiore, Guoqiang Yu, Jason J. Northey, Ronak Patel, Thomas A. Ravenscroft, Richard Ikegami, Wiert Kolkman, Pratik Kumar, Jonathan B. Grimm, Tanya L. Dilan, Virginia M.S. Ruetten, Misha B. Ahrens, Hari Shroff, Luke D. Lavis, Shaohe Wang, Valerie M. Weaver, Kayvon Pedram

**Affiliations:** Janelia Research Campus, Howard Hughes Medical Institute (HHMI), Ashburn, VA, USA; Center for Bioengineering and Tissue Regeneration, Department of Surgery, University of California, San Francisco (UCSF), San Francisco, CA, USA

## Abstract

All multicellular systems produce and dynamically regulate extracellular matrices (ECM) that play important roles in both biochemical and mechanical signaling. Though the spatial arrangement of these extracellular assemblies is critical to their biological functions, visualization of ECM structure is challenging, in part because the biomolecules that compose the ECM are difficult to fluorescently label individually and collectively. Here, we present a cell-impermeable small molecule fluorophore, termed Rhobo6, that turns on and red shifts upon reversible binding to glycans. Given that most ECM components are densely glycosylated, the dye enables wash-free visualization of ECM, in systems ranging from *in vitro* substrates to *in vivo* mouse mammary tumors. Relative to existing techniques, Rhobo6 provides a broad substrate profile, superior tissue penetration, nonperturbative labeling, and negligible photobleaching. This work establishes a straightforward method for imaging the distribution of ECM in live tissues and organisms, lowering barriers for investigation of extracellular biology.

## Introduction

The term “extracellular matrix” encompasses the cell membrane-tethered glycocalyx, the interstitial matrix that permeates the spaces between cells, basement membranes which provide a substrate for cell growth, and connective tissues such as fascia, tendons, and ligaments. The ECM is therefore a multi-scale and heterogeneous body-wide structure. Throughout the lifespan of an organism, extracellular matrices are actively remodeled by myriad cell types and play important roles in both biochemical and mechanical signaling^1,2^. For example, local increases in ECM stiffness can pattern the global orientation of developing mammary epithelium^3^, mechanical compaction of the ECM is sufficient to drive folding in engineered tissues^4^, and aberrant glycosylation can drive tumor immune evasion and metastasis^5^. In those examples, as well as many others, the three-dimensional arrangement of ECM biomolecules over time is critical to their individual activities and composite properties, motivating a longstanding interest in imaging the ECM in live tissues^6–8^.

Existing methods to fluorescently label extracellular biomolecules with affinity agents, genetic tags, and chemical labels, however, are challenging to apply in live tissues. Protein-based affinity reagents such as antibodies and lectins are severely limited by poor spatial diffusivity^9^. Genetic tagging of ECM components with fluorescent proteins can be challenging given viral packaging constraints and endogenous genetic tags require significant optimization to avoid perturbation of extracellular assemblies that are critical for developmental viability^10,11^. More broadly, antibodies and genetic tagging are typically used to visualize one or a few targets in a given sample, but are difficult to multiplex sufficiently to provide a comprehensive view of ECM structure, especially given heterogeneities in ECM composition across cells of a tissue, tissues of an organism, and organisms. These challenges are illustrated well by the dominance of label-free approaches such as second harmonic generation microscopy^12^ for visualization of the 28-member collagen family, as well as efforts to develop collagen-binding small molecule fluorophores^13,14^.

Glycosylation is a feature shared by nearly all ECM components^15^. As such, glycan-directed strategies have the potential to allow visualization of the ECM *en masse.* Glycan labeling techniques can be divided into two broad categories: metabolic incorporation with unnatural sugars and chemoenzymatic labeling^16^. Metabolic labeling is routinely used for imaging the glycocalyx in cultured cell systems and *ex vivo* tissue models, but generally requires >24 h of incubation with metabolic labels and is dependent on sample-specific glycosylation pathways^17^. Chemoenzymatic labeling, which is able to label a broader range of substrates, involves addition of an enzyme to the sample and is therefore subject to the spatial diffusivity of other protein-based methods^18^. An older technique for installing fluorophores on glycans, which involves sodium periodate oxidation followed by aniline-catalyzed oxime formation^19^, is toxic to live samples, necessitating staining at 4 °C and subsequent fixation. In general, since extracellular spaces, and by extension ECM, are not protected by a plasma membrane, methods that involve buffer exchanges are more likely to induce chemical, mechanical, and mass action driven perturbations to extracellular structures.

We envisioned that collective labeling of the ECM could be achieved with a cell-impermeable small molecule probe that increases in fluorescence upon reversible interaction with a chemical functionality found broadly on glycans (Fig. 1a). A small molecule would exhibit superior tissue penetration; low affinity, reversible binding would have the advantage of minimal perturbation to native structures and low photobleaching due to a large excess of unbound dye^20^. Further, by analogy to widely used DNA minor groove binding fluorogenic small molecules (e.g., Hoechst), such a dye could be employed as a one-step, wash-free dilution from a stock solution and would be applicable to a wide range of sample types. If successful, this method would lower barriers for testing hypotheses related to the composite properties of the ECM and by extension, to biological phenomena in extracellular spaces.

**Figure 1.**
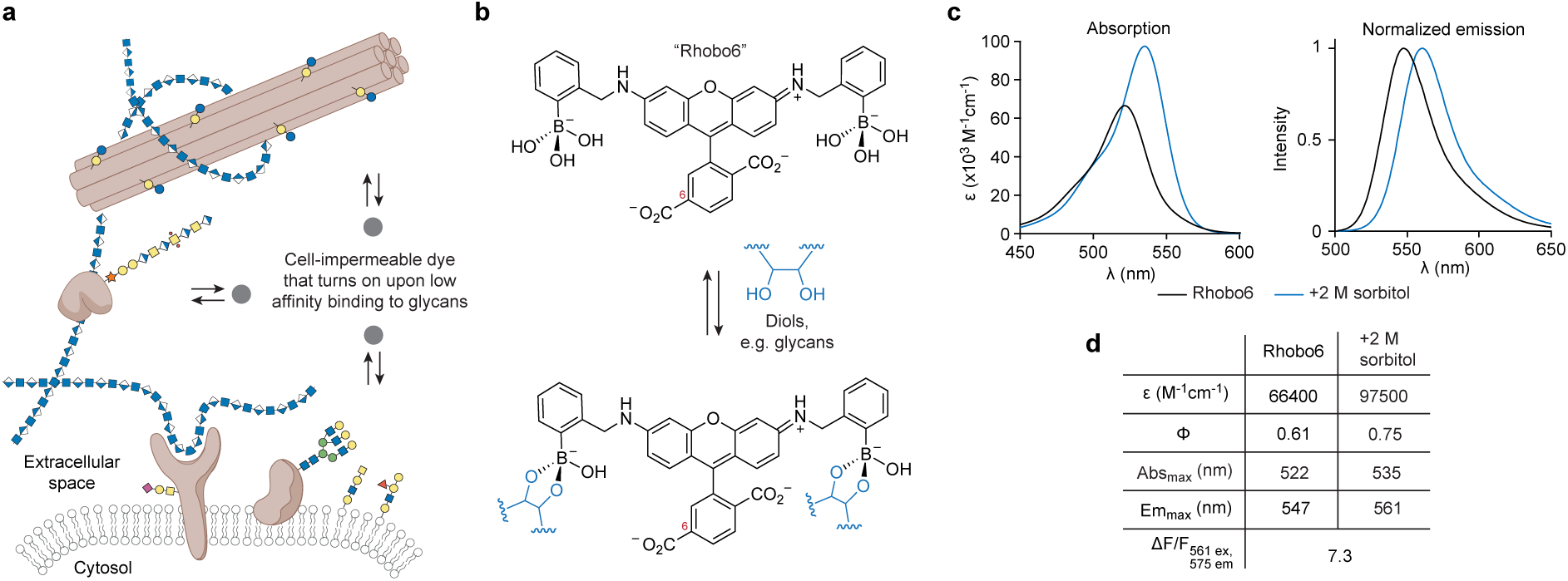
Photophysical characterization of the glycan-binding fluorophore Rhobo6. **a,** ECM labeling strategy. A cell impermeable dye is added to a biological sample such that it disperses into extracellular spaces. Upon reversible association with glycoconjugates (colored shapes) of the extracellular matrix, the dye increases its fluorescence output. **b,** Rhobo6 structure and propensity for glycan binding. The carboxylic acid on the 6-position of Rhobo6 (red numbering) is charged at physiological pH, rendering the molecule cell impermeable. The p*K*_a_ for *ortho*-aminomethylphenyl boronic acid is within the range of 5 to 7^30^, meaning the boronate and borate ester dominate in aqueous buffer at physiological pH. Rhobo6 is therefore expected to carry a net charge of negative three. **c,** Absorption and normalized emission spectra for Rhobo6 in unbound (5 µM dye in PBS) and bound (5 µM dye in PBS containing 2 M sorbitol) states. Emission spectra were measured with excitation wavelength at 490 nm. For 2P spectra see Extended Data Fig. 1i. **d,** Table of photophysical properties. Molar extinction (ε) is reported at peak absorption. Quantum yield (Φ) is measured as average value measured between 475 nm and 535 nm. Contrast is measured as relative fluorescence signal change between bound and unbound states (ΔF/F), when exciting at 561 nm and detecting fluorescence signal at 575 nm. Because of redshift in both absorption and emission, this value is highly dependent on both excitation and emission parameters (see also Extended Data Fig. 1e-h).

Boronic acids have been known for decades to exhibit reversible binding to 1,2- and 1,3- diols with dissociation constants in the tens of millimolar range^21^. Such diols are found in glycans and rarely elsewhere (e.g., on the ribose at the 3’ end of RNA), to the extent that boronic acids are employed for affinity purification of carbohydrates from complex biological samples^22^. In addition, boronic acids have a rich history of conjugation to fluorophores to give so-called “boronolectins”. This class of molecule, which includes boronated cyanines, rhodamines, BODIPY dyes, and others^23–25^, served as our starting point.

## Results

### Probe design and photophysical characterization

Of the previously described boronic acid-dye scaffolds, we were drawn to a Rhodamine 110- derived boronolectin, termed “Rhobo”, that had been developed by Strongin as a saccharide sensor for liquid chromatography^26,27^ and employed by Schepartz to bind tetraserine motifs on peptides^28^. First, Rhobo contains a phenylboronic acid on each side of the xanthene core, increasing affinity towards saccharides via avidity^29^. Second, Rhobo’s boronic acids are of the Wulff type, defined by the presence of an aminomethyl group *ortho* to the phenylboronic acid. Extensive studies by Anslyn, James, Shinkai, and Wang have shown that *ortho*-aminomethylphenyl boronic acids have the dual advantage of (i) lowering the p*K*a of the boronic acids and thereby enhancing the thermodynamics of sugar binding at neutral pH and (ii) increasing the kinetics of sugar binding via ammonium-mediated intramolecular general acid-catalysis^30^. Finally, fluorescence turn-on and spectral red shifts have been observed upon *in vitro* incubation of Rhobo with monosaccharides^27,28^.

Rhobo was reported to rapidly cross cell membranes and label intracellular structures when applied to cells at low micromolar concentrations, an observation that was confirmed with cultured cell monolayers (Extended Data Fig. 1a-b). Intracellular dye accumulation was unacceptable for imaging of ECM components, as it would (i) deplete dye from extracellular spaces and (ii) significantly reduce the signal to background ratio in cell-rich tissues.

Addition of functional groups that are charged at physiological pH is well known to reduce cell permeability of small molecule dyes (e.g., calcein^31,32^). A cell-impermeable Rhobo derivative was generated by addition of a carboxylic acid substituent at the 6-position of the dye, via a one-step reductive amination reaction with commercially available 6- carboxyrhodamine 110 and 2-formylphenylboronic acid (cf. *Methods*) (Fig. 1b). The resulting molecule, which we term “Rhobo6”, exhibited dramatically reduced cell permeability relative to Rhobo during 6 h of incubation on cultured cell monolayers (Extended Data Fig. 1a-b).

Photophysical characterization of Rhobo6 in diol-free buffer revealed an approximately 20- nm shift in absorbance and emission maxima relative to the parent dye 6-carboxyrhodamine 110, closely matching reported values for dibenzylrhodamine (i.e. a dye with *N*-benzyl groups lacking boronic acids)^33^. Next, a saturating concentration of the sugar alcohol sorbitol was added to generate the diol-bound form of the dye (Extended Data Fig. 1c-d). Relative to the unbound form, the bound form exhibited an increase in molar absorptivity, an increase in quantum yield, a 13 nm red shift in the absorbance peak, and a 14 nm red shift in the emission peak (Fig. 1c-d). Rhobo6 therefore turns on and red shifts upon binding diols. As a result of the red shift, the choice of excitation wavelength and emission filters will influence observed contrast (Extended Data Fig. 1e-h). Excitation with a 561 nm laser line coupled with a 575 nm longpass filter, corresponding to commonly used red fluorescent protein (RFP) imaging parameters, provided near-optimal fluorescence contrast, with a measured *in vitro* fluorescence change (ΔF/F) of 7.3. The two-photon (2P) excitation spectra of Rhobo6 exhibited an 800 nm peak, which increased in the diol-bound state (Extended Data Fig. 1i).

### Labeling profile for purified glycans and ECM components

To assess the specificity of Rhobo6 for glycans, a commercially printed glycan array was incubated with buffer containing 5 µM Rhobo6 and imaged without washing using a confocal microscope (Extended Data Fig. 2). Of the 100 glycans in the array, 98 showed a statistically significant increase in binding relative to negative controls, indicating a broad specificity for glycans and glycoconjugates. The glycans for which Rhobo6 showed relatively lower binding were enriched in negatively charged structures, suggesting that charge-charge interactions may influence Rhobo6 binding.

Next, Rhobo6 was applied at 5 µM in phosphate-buffered saline (PBS) to purified ECM constituents, including fibrillar glycoproteins (collagen I, fibronectin), network-forming glycoproteins (collagen IV, laminin), a proteoglycan (aggrecan), and a polysaccharide (hyaluronan) (Fig. 2a; images are not contrast normalized). Fluorescence contrast was observed across all substrates, with hyaluronan showing the weakest signal, possibly due to a lack of condensed structures (see discussion of K_D_, below). Pre-treatment with sodium periodate, which destroys 1,2-diols, reduced labeling of collagen I, laminin, and fibronectin, and pre-treatment with the glycosidase chondroitinase reduced labeling of aggrecan, indicating that Rhobo6-mediated fluorescence contrast is dependent on the presence of glycans (Fig. 2b and Extended Data Fig. 3a). Spectral imaging of a collagen I gel incubated with 5 µM Rhobo6 showed a red-shift in a region of interest (ROI) within the gel relative to an ROI within the buffer (Fig. 2c). Spatial mapping of the excitation maximum detected at each pixel gave a spectral contrast image (Fig. 2d). Comparison of spectral and intensity contrasted images confirmed the presence of both free and bound Rhobo6 in the field of view, with collagen-bound molecules exhibiting a red-shifted excitation maximum.

**Figure 2.**
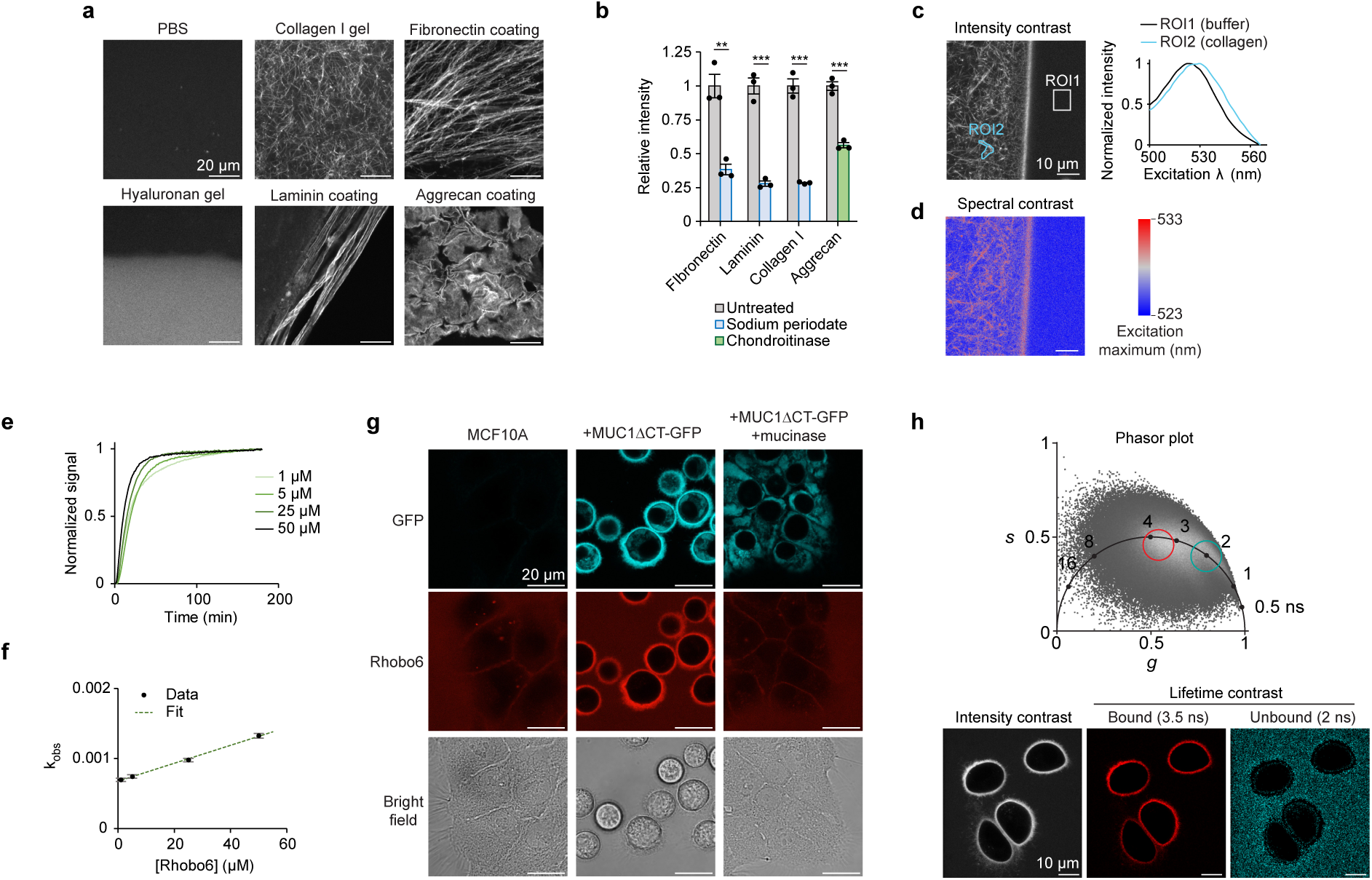
*In vitro* and *in cellulo* validation of Rhobo6 labeling. **a,** Rhobo6 labeling of purified ECM components. Substrates were prepared as glass coatings or gels (cf. *Methods*), and incubated with Rhobo6 at 5 µM for 1 h in PBS. Images were acquired with a confocal microscope. Contrast is not normalized across images. **b,** ECM components were treated with 10 mM sodium periodate (blue) or with Chondroitinase ABC (green), and signal intensity quantified from confocal microscopy images. For representative images used for quantification, see Extended Data Figure 3a. *N* = 3, error bars represent SEM. *P* values were determined by using two-tailed t-test; \*\**P* < 0.005; \*\*\**P* < 0.0005. **c,** Spectral imaging at the boundary of a collagen I gel and the surrounding buffer containing 5 µM Rhobo6, performed via excitation scan 500-566 nm and detection of fluorescence at 575-630 nm (cf. Supplementary Table 1). Intensity contrast image (*left*) obtained at 560 nm excitation with manually traced ROIs to capture an area rich in collagen fibers and an area within the surrounding buffer. Excitation spectra (*right*) corresponding to the manually drawn ROIs. **d,** A spectral contrast image generated by plotting excitation maxima for each pixel in (**c**). Binning = 2 pixels for the image shown. **e,** Time course of Rhobo6 fluorescence signal upon incubation with collagen I gels, at varying concentrations. Binding curves were used to extract a value for the observed equilibrium constant (k_obs_) at each concentration (cf. *Methods*). **f,** Linear fit between k_obs_ and Rhobo6 concentrations from (**e**), allowing extrapolation of binding constants k_on_ and k_off_. An apparent dissociation constant K_D_ of 53 µM was determined by the ratio of the two. Error bars represent 95% confidence interval for fitted k_obs_ values. **g,** Confocal microscopy of MCF10A cells labeled with Rhobo6. Expression of GFP-MUC1ΔCT was induced via addition of doxycycline. Mucin domains, which are N-terminal to GFP, were degraded enzymatically via live cell treatment with the mucinase StcE^5^. Note that mucin overexpression in these cells induces them to lift from their growth substrate, causing a spherical appearance with no loss in viability^35^. Contrast is normalized for each channel across experimental conditions. **h,** FLIM microscopy of MCF10A+GFP-MUC1ΔCT cells labeled with Rhobo6. Phasor plot (*top*) of lifetime distribution, with ROIs marking unbound and bound Rhobo6 population. *Bottom*, Intensity contrast image compared to lifetime bandpass images for each population (cf. Supplementary Table 1).

Next, to estimate an apparent dissociation constant (K_D_) for Rhobo6 binding to ECM substrates, the observed equilibrium constant (k_obs_) as a function of Rhobo6 concentration was measured using collagen I as a substrate (Fig. 2e)^34^. A linear fit allowed extrapolation of k_on_ (12.8 M^-1^s^-1^), k_off_ (6.77 x 10^-4^ s^-1^), and K_D_ (53 µM) (Fig. 2f). Our measured apparent dissociation constant is roughly two orders of magnitude greater than reported K_D_ values for binding of phenylboronic acid with monosaccharides^21^. These results suggest that a high effective molarity of diols is required for achieving substrate binding at micromolar concentrations of Rhobo6. At 5 µM Rhobo6 concentration, the time to reach fifty percent of maximum signal was 15 minutes, with over ninety percent of signal achieved at 60 mins, which motivated our incubation time of 1 h for biological samples.

Finally, the sensitivity of Rhobo6 labeling to photobleaching over repeated rounds of imaging was assessed. If, as expected, a reversible equilibrium existed between free and bound dye in a sample, the pool of excess free dye would replenish transiently bound photobleached molecules, resulting in a stable signal over time^20^. Indeed, using coated aggrecan as a substrate, no loss of fluorescence was observed over 9 h of acquisition at one frame per minute (Extended Data Fig. 3b-c and Supplementary Video 1).

### Glycan-dependent labeling on cell surfaces

Next, Rhobo6 was applied at 5 µM in serum-free media to an immortalized mammary epithelial cell line (MCF10A) in which the extent of cell surface glycosylation could be predictably modulated via doxycycline-inducible expression of the heavily *O*-glycosylated transmembrane protein Mucin-1 lacking its C-terminal cytosolic domain (MUC1ΔCT)^35^. After 1 h of incubation with Rhobo6 at 37 °C, MUC1-dependent Rhobo6 labeling of cell surfaces was observed, and this signal was ablated when cells were pre-treated with a mucin-selective protease (Fig. 2g)^5^. MUC1-dependent signal was reduced upon addition of exogenous sorbitol or serum-containing media, the latter of which is expected to be rich in glycoconjugates. Rhobo6 staining is not compatible with samples that are chemically fixed or otherwise exhibit compromised cellular membranes, as the dye will internalize, resulting in intracellular fluorescence that drowns out cell surface fluorescence signal (Extended Data Fig. 3d and Extended Data Fig. 4d).

Modulation of fluorescence lifetime upon changes of nitrogen atom substitution in rhodamine dyes has been reported^36^. Those results, alongside our observed increase in quantum yield upon diol binding, suggested that free and bound Rhobo6 populations could exhibit measurable differences in their fluorescence lifetimes. Indeed, fluorescence lifetime imaging microscopy (FLIM) of MUC1-expressing cells enabled gating of two populations, centered at 2 ns and 3.5 ns, which corresponded to free and bound dye populations, respectively (Fig. 2h).

### Benchmarking Rhobo6 in excised tissues

To further benchmark Rhobo6, samples with complex, multi-component extracellular matrices were required (Fig. 3a). We turned to mouse submandibular salivary glands isolated at embryonic day 13 or 14 (E13-E14) and cultured *ex vivo*. These glands continue to develop over the course of days in culture, undergoing budding and ductal morphogenesis^37^. To assess the biocompatibility of Rhobo6, growth and morphogenesis of paired salivary glands from seven embryos cultured with or without Rhobo6 over 48 h were assessed. Rhobo6 caused no difference in the overall morphology or the number of epithelial buds (Fig. 3b and Extended Data Fig. 4a), suggesting it is neither toxic nor perturbative to a primary embryonic organ explant.

**Figure 3.**
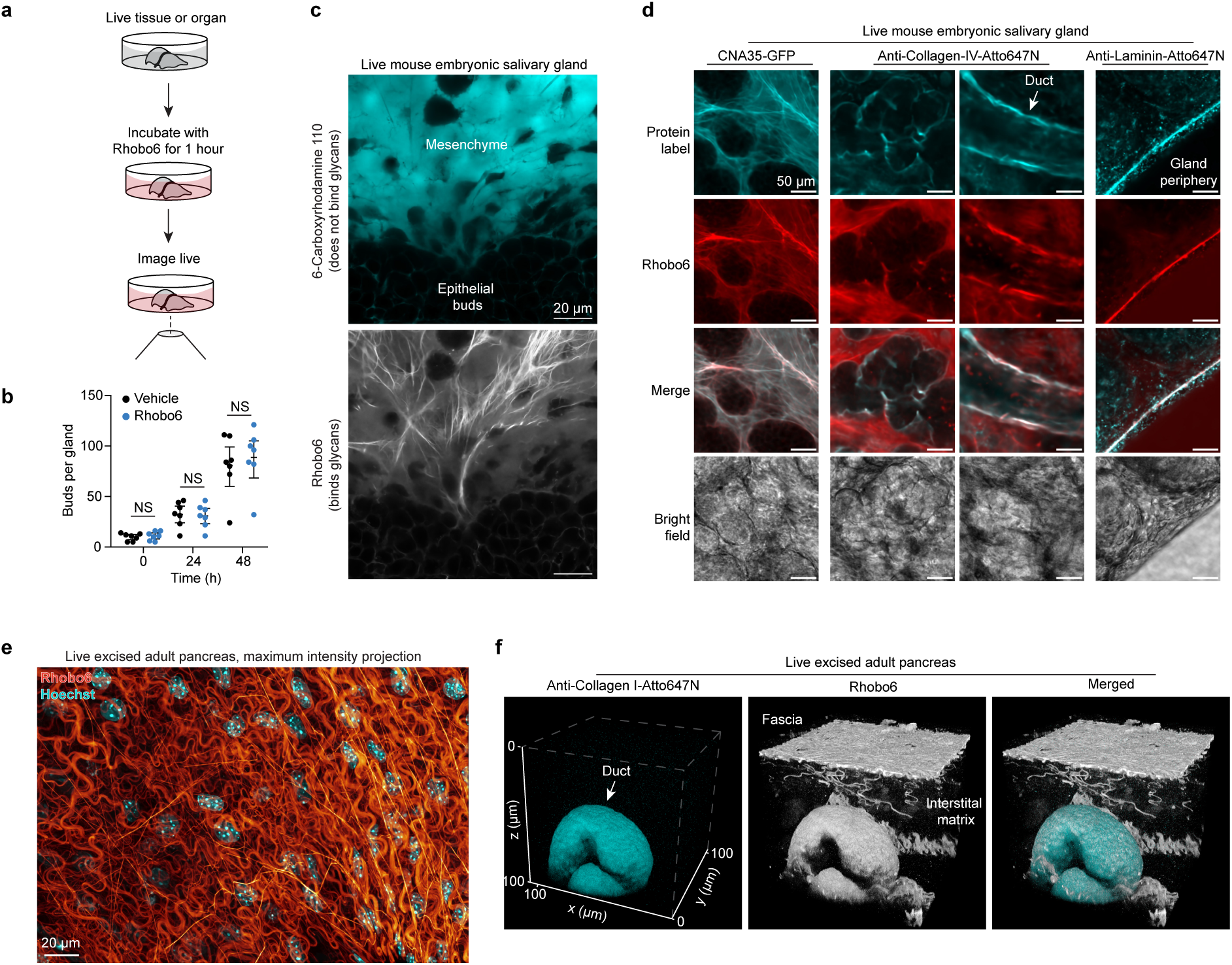
Labeling of excised tissues by bathing in Rhobo6-containing media. **a,** Cartoon illustrating the labeling approach. Freshly dissected or cultured tissues were labeled with 5 µM Rhobo6 for 1 h, with sample-specific media (cf. *Methods*). **b,** Viability and morphogenesis of mouse embryonic salivary glands upon incubation with Rhobo6. *N* = 14 glands were split into two paired groups, with each pair corresponding to glands from a single embryo. The first group was incubated with 5 µM Rhobo6 in media containing 0.5% DMSO, and the second group was incubated in media containing 0.5% DMSO, as a vehicle control. Viability and morphogenesis were assessed by counting epithelial buds every 24 h for 2 days. Paired groups were compared by paired t-tests; NS = not significant. **c,** Mouse embryonic submandibular salivary gland (E14) cultured *ex vivo* for 5 days, then labeled by bathing concurrently with Rhobo6 and 6-carboxyrhodamine 110. The latter dye differs from Rhobo6 only in that it does not contain the two *ortho*-aminomethylphenyl boronic acid groups, which are necessary for binding to extracellular glycans. Images were denoised (see Extended Data Fig. 4b-c for comparison of raw and denoised salivary gland images; Supplementary Table 1 reports image processing workflow for all datasets). **d,** Comparison of live Rhobo6 labeling to live labeling with protein-based affinity reagents against common ECM components, including fibrous collagen (CNA35), network-forming collagen (anti-collagen IV) and laminins. Glands were incubated with purified CNA35-GFP or Atto647N conjugated antibodies in solution along with Rhobo6, and imaged with a confocal microscope. Contrast not normalized. Images were denoised (cf. Supplementary Table 1). **e,** Freshly dissected and exsanguinated mouse pancreatic tissue, labeled by bathing with Rhobo6 (red), to highlight ECM, and Hoechst (cyan), to localize nuclei. Image shows a maximum intensity projection over a depth spanning 23 µm. Images were denoised (cf. Supplementary Table 1). **f,** Two-color labeling of exsanguinated adult mouse pancreatic tissue labeled by both Rhobo6 (red) and Anti-collagen-I-ATTO647N (cyan) antibody. Tissue was labeled by bathing for 1 h with both probes, and imaged with a confocal microscope. Image shows three-dimensional reconstruction of the (100 µm)^3^ volume.

Next, 6-carboxyrhodamine 110 and Rhobo6 each at 5 µM were added to a salivary gland for 1 h at 37 °C. Live two-color imaging was possible due to the ∼20 nm spectral separation of these two dyes (see above). The cell impermeable fluorophore 6-carboxyrhodamine 110, which does not contain boronic acids and therefore cannot bind glycans, filled extracellular spaces, similar to dextran-fluorophore conjugates and charged small molecule fluorophores that are routinely used for that purpose^31,32^ (Fig. 3c, *top*). Meanwhile, the boronic acid-functionalized dye Rhobo6, which binds to glycans, revealed a network of fibrillar material surrounding epithelial buds and mesenchymal cells (Fig. 3c, *bottom*). Next, Rhobo6 was compared with the previously reported cell-permeable analog Rhobo. Rhobo was not able to label structures of the ECM, likely due to depletion of the extracellular pool of dye following irreversible sequestration into epithelial and mesenchymal cells (Extended Data Fig. 4d-e).

To explore the identities of the molecules underlying observed Rhobo6 signal, glands were stained live with Rhobo6 and fluorescently labeled protein-based affinity reagents, followed by two-color imaging. In various fields of view, co-localization of Rhobo6 signal was observed with anti-collagen-IV antibody, with anti-laminin-1 antibody, and with CNA35, a 39 kDa truncation of the collagen adhesion protein from *S. aureus* which binds various forms of fibrillar collagen (Fig. 3d)^38^.

As a second test system, adult mouse pancreatic tissue was excised and bathed in buffer containing 5 µM Rhobo6 and 2 µg/mL Hoechst for 1 h. Confocal imaging of a 23 µm deep volume near to the tissue surface provided a view of the Rhobo6-stained fascia and embedded cellular nuclei, in a one-step, wash-free protocol (Fig. 3e). As the excised pancreatic tissue preparation is exsanguinated (cf. *Methods*), live antibody labeling was possible. The tissue was bathed with a fluorophore conjugated mouse anti-collagen I antibody alongside Rhobo6 for 1 h, then a 100 µm x 100 µm x 100 µm volume was acquired using a confocal microscope. The anti-collagen-1 antibody labeled a pancreatic duct, while Rhobo6 labeled the duct, surface fascia, and interstitial matrix (Fig. 3f), underscoring that Rhobo6 trades molecular specificity for a holistic view of ECM architecture. Since Rhobo6 is a rhodamine-based dye, it is compatible with live super-resolution imaging of the ECM using stimulated emission depletion (STED) microscopy^39^, as was demonstrated in freshly excised mouse pancreatic tissue (Extended Data Fig. 4f-h).

### Rhobo6 applied to non-mammalian model organisms

Glycosylation is a feature of extracellular biomolecules across the kingdoms of life. To test Rhobo6’s performance in non-mammalian systems, we turned to *D. melanogaster*, *C. elegans*, *D. rerio,* and *A. thaliana* model systems. In each case, our aim was to assess two features: (i) cell impermeability and (ii) ability to label material in the ECM.

Adult *D. melanogaster* brains were excised into saline containing 5 µM Rhobo6, incubated for 1 h at room temperature, then imaged live. A pattern of labeling was observed that suggested targeting of structures that surround neuronal cells, such as those in the mushroom body, central complex, and optic lobe (Extended Data Fig. 5a). Two-color imaging using a fly line with neurons expressing cytosolically targeted GFP confirmed that Rhobo6 labeling was excluded from cell interiors (Extended Data Fig. 5b). Adult *C. elegans* worms were injected with 10 pL of 100 µM Rhobo6 in each proximal arm of the gonad. Structures including yolk, eggshells, and the vulva were labeled (Extended Data Fig. 5c)^40^. A pattern of signal that based on bright field co-localization appeared to be within the oviduct was also observed. Two-color imaging with Rhobo6 in a worm line expressing endogenously tagged Nidogen-1-GFP^41^, however, revealed that the signal was concentrated at the sp-ut valve within the lumen of that cell, not within its cytosol (Extended Data Fig. 5d). In larval zebrafish (8 days post fertilization [d.p.f]), Rhobo6 was added at 5 µM to tank water and delivered via incisions to the tail. Rhobo6 visualized structural ECM components in the tail and notochord of the fish during a time-lapse of wound healing (Extended Data Fig. 5e and Supplementary Video 2). Finally, Arabidopsis seedlings were grown on agar from seed (cf. *Methods*). Seedlings were watered with 5 µM Rhobo6, incubated overnight, then imaged. Rhobo6 signal localized to root cell surfaces, consistent with previously observed distributions of metabolically incorporated azido-monosaccharides (Extended Data Fig. 5f)^42^. Taken together, these data confirm that Rhobo6 is compatible with a wide array of biological samples using a wash-free labeling protocol.

### Tissue distribution of Rhobo6 upon injection in mice

Given the absence of toxicity observed with application of Rhobo6 to developing salivary glands (Fig. 3b), we next investigated whether Rhobo6 could be administered to mice via injection (Fig. 4a). Retroorbital injection of 100 nmol (∼3.5 mg/kg) Rhobo6 did not result in apparent toxicity to 8-12 weeks old C57BL6/J females. To assess the distribution of the dye, mice were euthanized 30 mins post injection, and excised organs were placed on glass coverslips for imaging (cf. *Methods*). A panel of 12 live tissues was collected in this fashion using 2P microscopy, acquiring 2 mm x 2 mm areas and 70 µm x 70 µm x 50 µm volumes for each. Labeling of structures in the ECM was observed in all tissues except for the brain, where the dye is likely excluded by the blood brain barrier (Fig. 4b-d and Supplementary Videos 3-4; numbered arrows indicate tissue landmarks described in the caption to Fig. 4c-d). These images and volumes underscore the heterogeneity of ECM structures across tissues of the mouse and the broad distribution of Rhobo6 across organs, including relatively low blood flow areas such as tendon. Additionally, the presence of blood serum in these tissues did not interfere with Rhobo6 contrast, possibly due to the higher effective molarity of available diols in tissue ECM relative to cultured cell surfaces.

**Figure 4.**
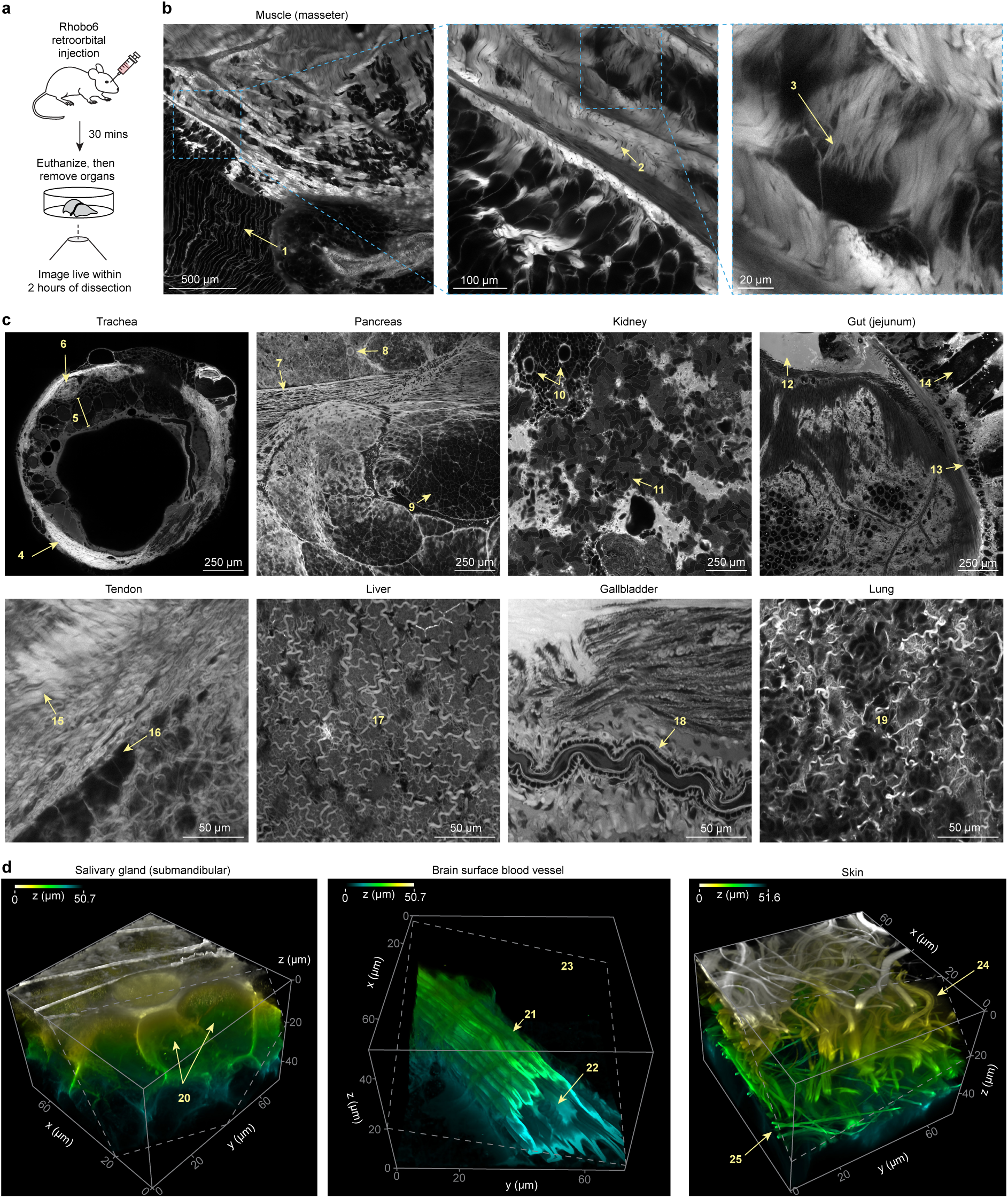
Rhobo6 distributes across mouse organs and labels the ECM upon retroorbital injection. **a,** Cartoon illustrating labeling approach. Anesthetized mice were injected retroorbitally with 100 µl of a 1 mM Rhobo6 solution in PBS containing 10% DMSO. Mice were allowed to recover for 30 min on a warming pad, then euthanized by cervical dislocation. Live tissues were harvested, placed on a glass bottom dish and imaged within 2 h of dissection. **b,** Two photon image of a 2 mm by 2 mm area of muscle tissue (masseter). Insets show sequential crops of the original image, highlighting both macroscopic and microscopic ECM features made visible by Rhobo6 labeling. For annotations of numbered landmarks see (**c**). **c,** Individual fields of view cropped from 2 mm by 2 mm two-photon images of the indicated tissues. Numbers in yellow correspond to features consistent with histological annotations^56,57^. Muscle (cf. (**b**)): (1) skeletal muscle fibers, (2) collagen-rich fascia, and (3) basal lamina surrounding myofibrils. Trachea: (4) tracheal cartilage ring, (5) submucosal layer with basement membrane, and (6) a tracheal gland encased in ECM. Pancreas: (7) longitudinal section of an interlobular duct, (8) the cross section of an intercalated duct, and (9) acinar tissue. Kidney: (10) a collecting tubule with a branching point, and (11) proximal and distal convoluted tubules. Jejunum: (12) mucus layer, (13) stratified squamous epithelial layer, and (14) villi. Tendon: (15) fascia of tertiary fiber bundle, and (16) fibroblasts. Liver: (17) entire field of view shows the fascia layer superficial to hepatocyte layer. Gallbladder: (18) longitudinal section of a capillary. Lung: (19) entire field of view shows alveolar tissue encased in ECM. All tissues, including images in (**b**) and (**d**), were acquired on the same day from 4 different mice of same strain and age (cf. *Methods*). Contrast not normalized across samples. **d,** Three-dimensional reconstructions of three tissues, from two-photon microscopy volumes. Depth color coding applied. Histological annotations are numbered in yellow. Salivary gland: (20) epithelial buds. Brain: (21) a blood vessel on the brain surface, (22) red blood cells excluded from Rhobo6 labeling within the vessel, and (23) brain tissue which is not labeled by Rhobo6, therefore appearing dark. Skin (24) collagen fibers and (25) elastin fibers. Contrast and depth-coded lookup table not normalized across samples. Images in (**d**) were denoised (cf. Supplementary Table 1).

For a subset of the tissues, head-to-head comparisons were performed with second harmonic generation microscopy and two-photon autofluorescence imaging, which are often used for imaging collagen and elastin, respectively^6^. Rhobo6 enabled visualization of both collagen and elastin structures simultaneously using between 15- and 40-fold lower light dose (1.1-3.3 J/cm^2^ vs 50.6 J/cm^2^ per sampled confocal voxel) along with a 40-fold lower detector gain (Extended Data Fig. 6).

### Intravital imaging of mouse mammary tumors

Malignancy in mammary tissue is accompanied by profound changes to the ECM, such as the accumulation and remodeling of fibrillar collagen into dense, linear and stiffened fibers^43–45^. To examine the potential for Rhobo6 labeling to help distinguish these ECM alterations at different stages of cancer progression, two photon intravital imaging was performed using mouse mammary tumor virus (MMTV)-driven polyoma middle T oncoprotein (PyMT) mice as representative of a well characterized genetically engineered mouse model of breast cancer.

PyMT rapidly induces spontaneous multifocal tumors in mice in a manner that is comparable to the stages of progression observed in human disease^46^. Mice were imaged at 10 weeks of age, when approximately 50% of MMTV-PyMT mammary glands contained advanced late carcinoma along with a mixture of adenoma/mammary intraepithelial neoplasia (MIN) and early carcinoma^46^. Two-photon imaging of tissue architecture in MMTV-PyMT mice was compared to that present in wild-type mammary glands (Fig. 5a). In live wild-type mice,

**Figure 5.**
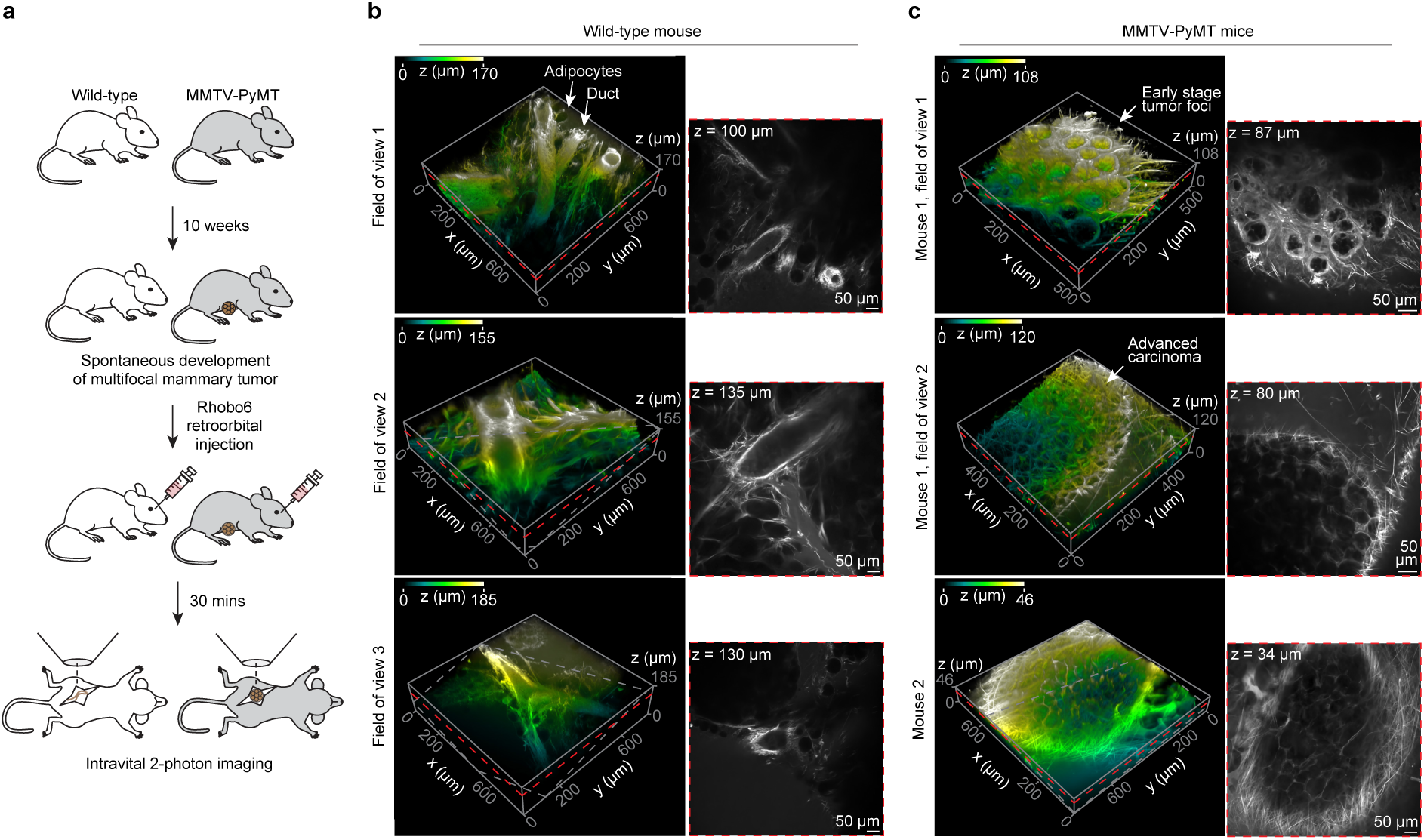
Intravital 2P imaging of ECM in a mouse model of mammary carcinoma. **a,** Cartoon representing experimental timeline, along with intravital imaging strategy for wild-type and mammary tumor bearing MMTV-PyMT mice. **b,** Rhobo6 imaging with three fields of view from the same mammary gland marking the extracellular matrix surrounding normal ductal architecture. *Left*, Volume rendering (depth-color coding applied). Arrows indicate adipocytes and epithelial ducts. *Right*, A single confocal slice from the adjacent volume (red dashed plane, with Z-height indicated) illustrating Rhobo6 labeling. Contrast is not normalized. Images were denoised (cf. Supplementary Table 1). **c,** The same as in (**b**) for two individual MMTV-PyMT mice. Two FOVs are presented for mouse 1 and one FOV for mouse 2. Arrows indicate early stage and late stage carcinomas. Contrast is not normalized. Images were denoised (cf. Supplementary Table 1).

Rhobo6 labeled the ECM surrounding ductal epithelium and fibrillar structures between stromal adipocytes (Fig. 5b). This contrasts with Rhobo6 imaging derived from MMTV-PyMT glands which demonstrated significant alterations to ECM architecture (Fig. 5c). Even at an adenoma/MIN or early stage of carcinoma (Fig. 5c, *top*), there was a thickening of the basement membrane around malignant foci and an increased presence of ECM between individual foci. In more advanced carcinoma, Rhobo6 distinguished a basket-like network of fibrillar ECM surrounding tumor nodules with many fibers oriented at more perpendicular angles rather than tangentially to the tumor margins; a phenotype associated with enhanced tumor cell invasion^47^.

Following intravital imaging, whole mammary glands were resected and fixed for histological analysis. Immunofluorescence labeling of actin (phalloidin) and cell nuclei (DAPI) was performed to visualize cellular architecture and localize malignant regions. In addition, CNA35 was included for comparison with Rhobo6-derived images. Differences in tissue architecture between mammary tumors and healthy ductal epithelium were apparent from the actin and nuclear labeling (Extended Data Fig. 7). Moreover, CNA35 mediated labeling demonstrated collagenous structures that corresponded well with the intravital imaging, with substantial infiltration of ECM and stroma between tumor foci and thick fibers oriented at increasing angles from the tumor margin (Extended Data Fig. 7). These studies confirm the ability of Rhobo6 to efficiently label ECM *in vivo* and effectively distinguish tumor-associated ECM from healthy ECM structures in an intravital imaging setting.

## Discussion

Our aim was to develop a method that enables one-step, wash-free visualization of ECM architecture in a wide variety of tissues. To achieve that goal, we developed a cell impermeable small molecule fluorophore, Rhobo6, that turns on and red shifts upon reversible binding to glycans, a nearly universal feature of ECM biomolecules. Rhobo6 has a number of characteristics that warrant discussion and will inform use cases.

First, a key enabling feature of Rhobo6 is its cell impermeability, which prevents irreversible intracellular accumulation and subsequent depletion of extracellular dye (Extended Data Fig. 4d-e). As such, Rhobo6 is incompatible with cellular samples that have been chemically fixed or where plasma membranes have been otherwise compromised (Extended Data Fig. 3d).

Second, the affinity of boronic acid groups for single monosaccarides is low, with dissociation constants expected to be in the range of tens of millimolar^21^. It is therefore expected that the high, local effective molarity of glycans in a biological sample drives the observed pattern of labeling (for estimation of effective K_D_ on a purified substrate see Fig. 2e-f). As a consequence, certain biological samples will have too low a density of available diols to be targeted by Rhobo6. For example, Rhobo6 appears to label the glycocalyx of cultured cells poorly (see MCF10A cells lacking MUC1ΔCT expression, Fig. 2g and Extended Data Fig. 3d), suggesting that methods such as metabolic incorporation and chemoenzymatic labeling^16^ will remain preferable to Rhobo6 for labeling the glycocalyx.

Third, when Rhobo6 is applied at low micromolar concentrations to a sample, an equilibrium exists between free and bound dye, with an excess of free dye available to replenish bound molecules. Such reversible, low affinity binding likely enables Rhobo6 to be minimally perturbative to native ECM structures in tissues, as demonstrated in the developing embryonic salivary gland (Fig. 3b and Extended Data Fig. 4a). The equilibrium also prevents photobleaching (Extended Data Fig. 3b-c and Supplementary Video 1), an advantage for studying tissue ECM dynamics which often occur over long timescales^1,2^. Finally, Rhobo6 labeling is reversible upon buffer exchange (Extended Data Fig. 4e), meaning repeated rounds of washing and labeling should enable spectral multiplexing and multi-timepoint imaging in a single sample^20^.

Fourth, Rhobo6 is designed to selectively bind extracellular glycans, but it does not exhibit specificity for any one glycan or ECM component (cf. glycan array, Extended Data Fig. 2). Rather, Rhobo6 broadly labels glycoconjugates of the ECM with a range of intensities that are not necessarily correlated with the abundance of the underlying biomolecules but instead the effective concentration and local environment of available diols. Imaging settings and image viewing settings therefore must be adjusted to highlight different ECM components (compare collagen I to hyaluronan in Fig. 2a and the laminin-rich band to collagen IV rich epithelial bud in Fig. 3d). Lowering image gamma, as was done for several tissues (cf. Supplementary Table 1), can overcome limitations in the dynamic range of viewing screens and human eyes.

Overall, Rhobo6 provides a holistic view of ECM architecture at the cost of molecular specificity. By analogy, live cell nuclear stains such as Hoechst take advantage of the fact that the nucleus is rich in a class of fundamental biopolymer (DNA) which has a unique motif (a minor groove). Though the specificity of Hoechst to DNA is complicated by base pair sequence preferences and by minor groove accessibility, in most sample types there is a sufficient quantity of substrate sites available for the dye to be used to visualize the distribution of nuclei in tissues^48^. Rhobo6, meanwhile, takes advantage of the fact that the ECM is rich in a different fundamental biopolymer (glycans) which also have a unique motif (1,2- and 1,3-diols). We envision Rhobo6 will find use as a straightforward and reliable counterstain for visualizing the distribution of ECM in tissues.

Looking ahead, opportunities exist for further development of phenylboronic acid modified fluorophores as ECM labels. First, a rich body of work suggests that the precise spacing of boronic acid groups can endow selectivity for monosaccharides and even oligosaccharides such as the Lewis antigen^24^. Second, though instability of Rhobo6 was not observed over days at room temperature (Extended Data Fig. 8b), enhanced oxidative stability may be advantageous in complex cellular environments and could be achieved with boralactones^21^. Finally, a color palette of Rhobo6 analogs could be generated via previously reported modifications to the rhodamine scaffold^27,49^. In all cases, molecular design efforts would be aided by characterization of the mechanism underlying both the fluorogenicity and spectral shift of Rhobo6 in the presence of sugars. Upon sugar binding, the majority of reported *ortho*-aminomethylphenyl boronic acid functionalized dyes exhibit turn on but do not red shift^30^. Rhobo6 differs from these molecules in that the *ortho*-aminomethyl group is directly attached via its nitrogen atom to the conjugated system of the fluorophore. Notably, a molecule synthesized by Shinkai in 1995 composed of an *ortho*-aminomethyl group attached in a similar fashion to the conjugated system of a coumarin also showed a spectral red shift upon sugar binding^50^.

Opportunities exist also for application of Rhobo6 using fluorescence imaging modalities aside from those presented here. Approaches for fast volumetric imaging such as two-photon structured illumination microscopy (2P ISIM)^51^ and adaptive optics lattice light sheet microcopy (AO-LLS)^52^ and could be applied to visualize ECM dynamics within scattering and inhomogeneous tissues. A low concentration of Rhobo6 applied to relatively immobile samples may allow points accumulation for imaging in nanoscale topography (PAINT) microscopy, which provides nanometer precision single molecule localizations^53,54^. Generation of a large ground truth dataset of known labels co-localized with Rhobo6 may open the door to machine vision annotation of ECM components based on properties such as persistence length and cellular context, providing a degree of molecular information in single-color Rhobo6 images^55^.

Finally, our intravital imaging results suggest that Rhobo6 or future analogs may find utility as diagnostic tools for human biopsy samples, in diagnostic imaging, or in fluorescence-guided surgery. To explore that possibility, dye pharmacokinetics will need to be characterized and fluorescence contrast in clinical settings will need to be assessed.

## Acknowledgements

We thank Andrew Krasley, Daniel Feliciano, Boaz Mohar, Damien Alcor, Ellen Quarles, Isabel Espinosa-Medina, Allyson Sgro, Aishwarya Korgaonkar, Jacob Smith, Christopher Obara, and William Lemon (all at Janelia), along with Stephen Adams (University of California, San Diego), Thomas Boltje (Radboud University), and Helene Langevin (NIH) for helpful discussions. We thank Gerald Rubin, David Clapham, and Timothy Brown (all at Janelia) for helpful discussions and for providing support to T.A.R., T.L.D., and R.P., respectively, during the course of this work. We thank Meng Wang and Tao Chen (Janelia) for advice on two-photon and second harmonic generation microscopy. We thank Andrian Gutu (Janelia) for access to a 2P microscope. We thank Matthew Paszek (Cornell) for MCF10A GFP-MUC1ΔCT cells. We thank the staff within the Biological Imaging Development CoLab (BIDC) at UCSF Parnassus Heights for training and use of 2P and spinning disk microscopes. *C. elegans* strains were provided by the Caenorhabditis Genetics Center, which is funded by NIH Office of Research Infrastructure Programs (P40 OD010440). Work at UCSF was supported by National Institutes of Health grant 1R35CA242447-01A1 to V.M.W. Work at Janelia was supported by the Howard Hughes Medical Institute.

## Author contributions

A.F. and K.P. designed, performed, and analyzed experiments unless noted otherwise. G.Y. and W.K. performed chemical syntheses, with advice from P.K., J.B.G., and L.D.L. P.K. designed the Rhobo6 stability experiment. J.J.N. and V.M.W. designed and performed intravital mouse tumor experiments. R.P. and L.D.L. contributed to photophysical characterizations. T.A.R. contributed to *D. melanogaster* experiments. R.I. and H.S. contributed to *C. elegans* experiments. V.M.S.R. and M.B.A. contributed to *D. rerio* experiments. T.L.D. contributed to excised pancreas experiments. S.W. contributed to salivary gland experiments and designed and analyzed the gland viability time course experiment. A.F. and K.P. wrote the manuscript with input from all authors.

## Methods

### Microscopy methods

Supplementary Table 1 tabulates microscopy platforms, imaging parameters, and data processing steps for all datasets. Unless noted otherwise, image processing was performed in Fiji/ImageJ (National Institute of Health, USA).

#### Considerations for multiplexing

Rhobo6, particularly in the unbound state, can be excited by a 488 nm laser line to some degree (see Extended Data Fig. 1e-h). As a result, multiplexing with green fluorophores, e.g. GFP, requires attention to emission filters to minimize fluorescence crosstalk. In particular, we typically employed a cut-off wavelength of 525 nm for the green emission filter, while keeping the Rhobo6 emission filter above 575 nm (Supplementary Table 1). For multiplexing with far red probes such as Atto647N, we set the upper cutoff for the Rhobo6 emission filter to 630 nm.

### Organic synthesis and chemical characterization

#### General considerations

All chemicals were obtained from commercial suppliers in reagent grade or higher and used as received. Reactions were conducted in 2-5 mL Biotage microwave vials sealed with Biotage microwave proof caps and heated in a Biotage Initiator+ microwave synthesizer. Reactions were monitored by LC/MS (Phenomenex Kinetex 2.1 × 30 mm 2.6 µm C18 column; 2-10 µL injection; 5-98% MeCN/H_2_O, linear gradient, with constant 0.1% v/v HCO_2_H additive; 6 min run; 0.5 mL/min flow; ESI; positive ion mode). Reaction products were purified by preparative HPLC (Phenomenex Gemini 30 × 150 mm 5 µm NX-C18 column). Analytical HPLC analysis was performed with an Agilent Eclipse 4.6 × 150 mm 5 μm XDB-C18 column under the indicated conditions. High-resolution mass spectrometry was performed by the High Resolution Mass Spectrometry Facility at the University of Iowa. NMR spectra were recorded on a Bruker Avance II 400 MHz spectrometer. Chemical shifts are reported in parts per million (ppm) relative to residual solvent peaks. ^1^H NMR data are presented as follows: chemical shift (δ ppm), multiplicity (s = singlet, d = doublet, t = triplet, q = quartet, dd = doublet of doublets, dt = doublet of triplets, dtd = doublet of triplet of doublets, m = multiplet), coupling constant in Hertz (Hz), integration.

(E)-2-(6-((2-boronobenzyl)amino)-3-((2-boronobenzyl)iminio)-3H-xanthen-9-yl)benzoate (Rhobo) was synthesized as previously reported^58^.

#### Synthesis of (E)-2-(6-((2-boronobenzyl)amino)-3-((2-boronobenzyl)iminio)-3H-xanthen-9-yl)-4-carboxybenzoate (Rhobo6)

6-Carboxyrhodamine 110 (HCl salt, 51 mg, 0.124 mmol, 1 eq), 2-formylphenylboronic acid (100 mg, 0.667 mmol, 5.4 eq), sodium triacetoxyborohydride (90 mg, 0.425 mmol, 3.4 eq), and anhydrous DMF (1.7 mL) were added to a microwave vial containing a magnetic stir bar. The mixture was homogenized by ultrasonication at room temperature for 1 min, and concentrated acetic acid (50 μL, 0.874 mmol, 7.0 eq) was added. The vial was sealed with a microwave proof cap and stirred at room temperature for 1 min. The reaction was then run at 130 °C for 60 min in the microwave synthesizer. After the vial was cooled to room temperature, the cap was removed and ∼20 mL of MeOH was added to dilute the reaction mixture. Purification by preparative HPLC (25 - 50% MeCN/H_2_O, linear gradient, with constant 0.1% v/v TFA additive) yielded red Rhobo6 solid (TFA salt, 33.2 mg, 35%). ^1^H NMR (400 MHz, CD_3_OD): δ 8.40 – 8.36 (m, 2H), 7.97 –7.95 (m, 1H), 7.42 – 7.28 (m, 8H), 7.04 (d, *J* = 9.4 Hz, 2H), 6.89 (dd, *J* = 9.3, 2.2 Hz, 2H), 6.79 (s, 2H), 4.73-4.60 (m, 4H). Analytical HPLC: t_R_ = 10.4 min, 95.4% purity (10 – 95% MeCN/H_2_O, linear gradient, with constant 0.1% v/v TFA additive; 20 min run; 1 mL/min flow; ESI; positive ion mode; detection at 254 nm); HRMS (ESI) calculated for C_35_H_28_B_2_N_2_O_9_ [M+H]^+^ 643.2054, found 643.2065.

#### Dye storage

Freshly prepared Rhobo6 solid was dissolved in anhydrous DMSO at 10 mM, and subsequently distributed into 10 µl aliquots in screw top vials, which were stored at −80 °C. Unless noted, aliquots were thawed at room temperature, diluted with anhydrous DMSO to 1 mM then frozen once again at −80 °C. 1 mM DMSO aliquots were diluted 1:200 into sample buffer to yield 5 µM working concentrations, and freeze/thawed 5 times or fewer before being discarded.

#### Stability of Rhobo6 by HPLC

An aliquot of Rhobo6 (10 µl of 10 mM in DMSO) was diluted with DMSO (40 µL) and PBS (50 µL) to a final concentration of 1 mM and then stored at room temperature in darkness. Its purity was assessed at 0, 1, 2, 4, 8, 24, 48, and 72 h by an analytical HPLC (Phenomenex Gemini 4.6 × 250 mm 5 µm NX-C18 column; 5 µL injection; 5 – 95% MeCN/H_2_O, linear gradient, with constant 0.1% v/v TFA additive; 25 min run; 1 mL/min flow; detection at 254 nm).

### Monosaccharides and monosaccharide analogs

0.4 M solutions of D-Glucose (Millipore Sigma, G7021), D-Galactose (Fisher Scientific, BP656-500), D-Mannose (Millipore Sigma, M8574), D-Fructose (Millipore Sigma, F0127), sorbitol (Millipore Sigma, PHR1006) were prepared in 0.5 mL PBS. The pH of all solutions was adjusted to be in the 7.3-7.4 range using a pH-meter (Mettler Toledo S470 coupled with Ultra-Micro-ISM pH probe) before reaching the total PBS volume. Three 10 µM Rhobo6 solutions were prepared separately in 10 mL of PBS starting from three different 10 mM aliquots of dye. Each compound was measured in triplicates, and for each of the three measurements 25 µL of 0.4 M monosaccharide solution was combined with 25 µL of one of the dye solutions. Dilution was performed directly in a 96 well plate with black walls (Greiner Bio-One, 655900), followed by 1 h incubation to allow for binding to stabilize. Fluorescence emission was then measured using a Tecan Spark plate reader, with 555 nm excitation and 570 to 630 nm emission range with 2 nm step, and 40 µs integration time. As controls, both PBS (buffer only) and each of the dye solutions (dye only) were also acquired. Spectra were then analyzed by subtracting background counts (estimated by the buffer only signal at each wavelength) and averaging the maximum signal for each spectrum across the three measurements.

### Photophysical characterization

Rhobo6 solutions were prepared by diluting 1 mM dye stock in either PBS (Corning, MT21040CV) or PBS solution containing 2 M sorbitol (Millipore Sigma, S1876). Absorbance measurements were performed using a UV-Vis spectrometer (Cary 100, Agilent technologies) at 5 µM dye. Extinction coefficients were calculated at peak absorbance in both conditions. Fluorescence emission spectra were measured using a spectrofluorometer (Cary Eclipse, Varian Inc.) with excitation set at 490 nm. Quantum Yield measurements were performed using an integration sphere spectrometer (Quantaurus, Hamamatsu), averaging values measured between 475 and 535 nm, with a 5 nm increment.

To measure the contrast at different excitation wavelengths, we acquired emission spectra of both bound and unbound solutions while exciting at 490 nm and 561 nm. The reported contrast ΔF/F_561ex/575em_, i.e. the fluorescent contrast at 561 nm excitation and 575 nm emission, was calculated as relative signal change at 575 nm emission, calculated from the 561 nm excitation dataset.

To renormalize spectra to a given excitation wavelength both emission spectra were normalized to a wavelength-specific excitation coefficient. This coefficient was estimated as:

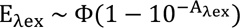

Where E_λex_ denotes the estimated excitation at a given excitation wavelength λ, Φ denotes the measured quantum yield, and A_λex_ denotes the measured absorbance at λ.

#### Two photon excitation

Two-photon excitation spectral measurements were performed as previously described^59^. Dye solutions of 1 µM concentration in 100 mM phosphate buffer, pH 7.4 or the same with 1 M galactose were prepared. Spectral measurements were performed using an inverted microscope (IX81, Olympus) equipped with a 60X, 1.2NA water immersion objective (Olympus). Dye samples were excited with pulses from an 80 MHz Ti-Sapphire laser (Chameleon Ultra II, Coherent) for 710-1080 nm range, and with an OPO (Chameleon Compact OPO, Coherent) for the 1000-1500nm range. Fluorescence collected by the objective was filtered through a dichroic (675DCSXR, Omega and FF825-SDio1, Semrock) and an emission filter (539BP278 and 709BP167, Semrock), before detection by a fiber-coupled Avalanche Photodiode (APD) (SPCM_AQRH-14, Perkin Elmer). Two-photon excitation spectra were obtained from 1 µM dye samples at 1 mW of laser power across the spectral range of 710 nm to 1080 nm using Ti:Saphire and at 2 mW of laser power for spectral range of 1000-1500 nm using OPO. The excitation spectra have been normalized for the laser power and corrected for the transmission of the dichroic, emission filter, and quantum efficiency of the detector across wavelengths. As control, we acquired and reported RhodamineB two-photon excitation spectrum measured in the same manner, which is also reported in the literature^60^. All spectra are averages of *n* = 2 measures.

### Glycan array

To test specificity of glycan binding across various glycoconjugates we used a glycan array (RayBiotech, GA-Glycan-100-1), which is manufactured as a glass slide divided in wells containing one array replicate; each array consists of 100 different glycans printed in 4 replicate spots, along with two sets of 4 negative control spots. The glass slide was equilibrated from storage temperature (−20 °C) to room temperature for 90 minutes; subsequently wells were rehydrated by incubating in PBS for 60 minutes. Then, PBS buffer was replaced with a 5 µM Rhobo6 solution in PBS. Three arrays were incubated at least 90 minutes before being imaged sequentially with a confocal microscope. From incubation to imaging completion a maximum of 6 h elapsed. Fluorescence signal and local background for each glycan was quantified using a MATLAB (R2022a, MathWorks) script receiving user input, followed by a background corrected signal normalization within each array to account for slight changes in imaging conditions. Data was visualized and analyzed for statistical significance using PRISM (GraphPad); statistical significance was determined through Dunnett-corrected t-test for multiple comparisons to a control group, with assumption of unequal variance across groups.

### Purified extracellular matrix components

#### Coatings

Human fibronectin (Corning, 354008) was resuspended in water at 1 mg/ml, gently mixed and incubated at 37 °C for 20 minutes before being used or aliquoted and frozen at −20 °C. Laminin (Thermo Fisher Scientific, 23017015) is supplied at 0.5-2 mg/ml in 50 mM Tris-HCL (pH 7.4), 0.15 M NaCl and was aliquoted and stored at −20 °C. Lyophilized aggrecan (Millipore Sigma, A1960-1MG) was resuspended at 2 mg/ml in PBS, aliquoted and stored at −20 °C. After thawing an aliquot of each solution at 4 °C, 10 µl of each substrate was deposited in an untreated glass-bottom well (Ibidi, 80807), and dried over a hot plate set to 37 °C for 4 h prior to use.

#### Collagen gels

Collagen I gels at 0.15 mg/ml were prepared starting from collagen type I (Ibidi, 50201) thawed at 4 °C over few hours and diluted in 17.5 mM acetic acid to 4 mg/ml. A solution was then prepared with 6.67% 10x DMEM (Millipore Sigma, D2429-100ML), 6.67% NaOH 1M solution in water, 18.5% distilled water, 3.33% of NaHCO_3_ 89 mM solution in water, 33.33% of 1x DMEM (Thermo Fisher Scientific, 21041025) and 37.5% of collagen I solution at 4 mg/ml. The solution was quickly distributed in 20 µl droplet in an untreated glass-bottom well (Ibidi, 80807), and incubated at 37 °C, 95% humidity, 5% CO_2_ for at least one hour prior to use.

#### Hyaluronan gels

To high MW hyaluronan (1500 kDa, 10 mg, R&D Systems) in a 1.5 mL Eppendorf microcentrifuge tube was added 90 μL of 0.25 M NaOH aqueous solution. The mixture was centrifuged at 5500 rpm and vortexed for 2 min, respectively, until the hyaluronan was dissolved. 1,4-Butanediol diglycidyl ether (BDDE, 1 μL) was diluted with 0.25 M NaOH aqueous solution (10 μL) and then added to the hyaluronan solution. The mixture was centrifuged at 5500 rpm and vortexed for 30 s, respectively, and then centrifuged at 22500 rpm for 5 min to get rid of bubbles. 2 μL of the solution was pipetted onto each well of an 8 well plate. The well plate was surrounded with water to prevent the hyaluronan solution from drying and placed in an oven at 40 °C for 16 h during which time the hyaluronan was crosslinked by the BDDE to form a gel.

#### Sodium periodate and chondroitinase treatments

A 10 mM solution of sodium (meta)periodate (MilliporeSigma, 71859-100G) was prepared in PBS, added to substrate wells and incubated for 6 h. The reaction was then quenched with 0.1 M glycerol solution. At this point the wells were washed three times with PBS and subsequently imaged. ChondroitinaseABC (MilliporeSigma, C3667-10UN) was aliquoted at 50 units/mL in PBS. Upon use, aliquots were diluted 1:15 in PBS, added to the treated wells. Samples were then incubated for 6 h, and subsequently washed three times with PBS and imaged. Coatings and gels were prepared in triplicates for both untreated and treated conditions. Collagen-I, fibronectin and laminin were treated with sodium periodate while aggrecan was treated with ChondroitinaseABC. Three control wells were left with PBS only as control. After treatment, all wells were incubated in a 5 µM Rhobo6 solution in PBS for 1 h and imaged with a confocal microscope. Fluorescence signal was quantified in each field of view as average intensity in a manually traced region of interest containing the signal.

### Spectral imaging

A collagen-I gel was prepared following methods described above, and then incubated in a 5 µM Rhobo6 solution in PBS for 1 h. Excitation scan modality was selected on a Leica Stellaris 8 confocal microscope, scanning excitation wavelength from 500-566 nm with 2 nm incremental steps, and emission detected in the 575-630 nm range. The result was a three-dimensional image in which each pixel contains an absorption spectrum. Spectra for ‘Collagen’ and ‘PBS’ was plotted averaging pixel value in manually traced regions of interest. Spectral contrast image was generated plotting the maximum absorbance wavelength in each pixel using MATLAB (R2022a, MathWorks).

### Estimation of Rhobo6 binding affinity

Binding affinity to collagen type-I was measured following the fit of equilibrium constants^34^. Four collagen-I gels (10 µL volume each) were prepared following methods described above in a 50 mm glass bottom petri dish (MatTek, P50G-1.5-14-F). Right after deposition of collagen solution on glass, a gentle tap was applied to distribute the gel on a larger surface, reducing its thickness. Upon use, each gel equilibrated at room temperature in 2 mL of PBS for 60 minutes. Then, it was placed on a confocal microscope and a focal plane within the collagen gel was established using brightfield contrast. The microscope was set up to acquire a 3 h timelapse at 1 frame per minute rate. After the first timepoint was acquired setting initial conditions, 2 mL of a Rhobo6 solution at twice the target concentration were added, resulting in a 1:1 dilution. The concentration of Rhobo6 solution was changed each time as the independent variable. After the acquisition, the mean intensity over the full field of view was extracted, and the observed equilibrium constant k_obs_ determined by fitting the following equation in MATLAB (R2022a, MathWorks):

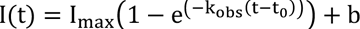

Where *I* is the measured intensity, *t* is the time, *b* is the background intensity and *t_0_* is a time delay parameter to take into account the arbitrary moment in which dye solution was added.

Once all binding curves were acquired, a linear fit was performed between concentration and observed equilibrium constant following the relationship:

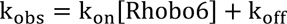

Where k_on_ and k_off_ are the binding and unbinding constants, respectively. A dissociation constant K_D_ for collagen type-I was thus estimated as K_D_=k_off_/k_on_.

### Photobleaching test

A well within a glass bottom 8-well plate (Ibidi, 80807) was coated with aggrecan following the protocol described above and incubated overnight with a 5 µM solution of Rhobo6 in PBS. The well was then completely filled with Rhobo6 solution and sealed with parafilm to prevent evaporation. The sample was imaged with a confocal microscope at 1 image per minute overnight. The signal was plotted over time, calculated as the mean pixel value in manually traced ROIs containing the sample.

### Mammalian cell culture experiments

#### Cell culture conditions

Cells were maintained at 37 °C and 5% CO_2_. MCF10A GFP-MUC1ΔCT cells (Paszek Lab, Cornell) were cultured in phenol red-free 1:1 DMEM:F12 supplemented with 5% New Zealand horse serum (Thermo Fisher Scientific, 16050122), 20 ng/ml epidermal growth factor (Peprotech), 0.5 μg/ml hydrocortisone (Millipore Sigma, H0888-1G), 100 ng/ml cholera toxin (Millipore Sigma, C8052-.5MG), 10 μg/ml insulin (Millipore Sigma, I1882-100MG) and 1% penicillin/streptomycin (P/S) (Thermo Fisher Scientific, 15070063). PC-3 (ATCC, CRL-1435) cells were cultured in RPMI-1640 (Thermo Fisher Scientific,11875093) supplemented with 10% heat-inactivated fetal bovine serum (Thermo Fisher Scientific, 10-438-026) and 1% P/S.

#### Dye penetration experiment

PC3 cells were plated in an 8-well plate (Ibidi, 80807) at 10,000 cells per well and cultured for 2 days. Rhobo or Rhobo6 were added at 5 µM via a 1:200 dilution from a 1 mM DMSO stock. Images were acquired on a confocal microscope upon addition (t =0) and after 1, 2 and 6 h; cells were incubated at 37 °C and 5% CO_2_ between time points. Intracellular dye presence was quantified as average fluorescence signal within manually traced ROIs along cell perimeters for *N* = 9 cells for each condition and timepoint.

#### MCF10A GFP-MUC1ΔCT experiments

Cells were plated at 10,000 cells per well in an 8-well dish (Ibidi, 80807) and cultured for 2 days. Doxycycline (Panreac AppliChem, A2951) was added a 1 µg/mL via a 1:1000 dilution from a 1 mg/mL stock in DMSO and cells were incubated for another 2 days. Doxycyline induces the overexpression of the surface glycoprotein MUC1 that is lacking its C-terminal cytosolic domain (MUC1ΔCT).

Overexpression of MUC1ΔCT causes cells to ball up and lift from their growth substrate^35^. Once lifting was observed in the majority of cells, the wells were washed 1x with PBS, and the PBS was replaced with the indicated media containing 5 µM Rhobo6. Fixation was performed with 4% paraformaldehyde (Electron Microscopy Sciences, 19202) in PBS for 30 mins at room temperature and mucinase treatment was performed with 100 nM StcE mucinase (expressed and purified as previously reported^5^) for 4 h at 37 °C.

### FLIM microscopy

MCF10A cells expressing GFP-MUCΔCT were plated and cultured as described above. Before imaging, cells were washed 3 times in PBS, and then incubated in a 5 µM Rhobo6 solution in PBS for 1 h at standard cell culture conditions (37 °C, 95% humidity, 5% CO_2_). FLIM microscopy was performed using an Abberior Facility Line microscope. Lifetime contrast images, phasor plot coordinates and lifetime bandpass images were generated within the microscope software. The reported phasor plot was generated in MATLAB (R2022a, Mathworks).

### Mouse embryonic salivary glands

#### Dissection and culture

All experiments complied with protocols approved by the Institutional Animal Care and Use Committee (IACUC) at Janelia Research Campus (protocol number 22-0230). Submandibular salivary glands (SMG) were harvested and cultured *in vitro* following the protocol reported by Wang *et al.*^37^. Briefly, mouse submandibular salivary glands were dissected from 13- to 14-day old embryos (E13-E14). The embryos were isolated from timed pregnant CD-1 outbred mice (Charles River Laboratories). Isolated salivary glands were cultured on 13 mm diameter 0.1 µm pore polycarbonate filters (MilliporeSigma, WHA110405) floating on 1 mL Organ Culture Medium (see below) in a 35-mm dish at 37°C with 5% CO2. Base Medium was DMEM/F-12 (Thermo Fisher Scientific, 11039047) supplemented with 2 mM L-glutamine (Thermo Fisher Scientific, 25030081) and 1x PenStrep (100 units/mL penicillin, 100 µg/mL streptomycin; Thermo Fisher Scientific, 15140163). Organ Culture Medium was Base Medium supplemented with 150 µg/mL vitamin C (MilliporeSigma, A7506) and 50 µg/mL transferrin (MilliporeSigma, T8158).

#### Rhobo6 incubation and mounting for imaging

For Rhobo or Rhobo6 dye labeling, 5 µL of 1 mM stock was diluted in 1 mL of Organ Culture Medium to make a 5 µM labeling solution. Culture medium was replaced with the labeling solution followed by a 1 h incubation at 37 °C with 5% CO_2_. In order to mount the samples for inverted microscope imaging, we used double-adhesive imaging spacers (Grace Bio-labs, 654002) attached to the 27 mm glass wide bottoms of 35 mm dishes (Thermo Fisher Scientific, 150682). Under a dissecting microscope, 5 µL Organ Culture Medium was transferred to the center of the imaging spacer, and the filter with glands was flipped onto the imaging spacer using a pair of forceps so that glands were sandwiched between the filter and the dish bottom. Care was taken to ensure the filter was flat and center-aligned with the imaging spacer. The edge of the filter was pressed to ensure tight adherence to the imaging spacer. 1 mL Organ Culture Medium with 5 µM Rhobo dye was then added on top of the flipped filter. The mounted glands were imaged right away.

#### Toxicity test during ex vivo culture

To evaluate whether Rhobo6 adversely affects the growth or branching morphogenesis of *ex vivo* cultured embryonic salivary glands, paired glands from the same embryo were separated into two groups, which were treated with 5 µM Rhobo6 (1 mM stock in DMSO) or 0.5% DMSO. Phase contrast images of cultured glands were acquired at 0, 24, and 48 h. The number of buds were counted manually on these images in Fiji using an ImageJ macro to facilitate recording of the results. To minimize bias, file names of all images were scrambled for observer blinding using a Python script before counting. The counting results were subsequently decoded and analyzed using customized Python scripts. The paired t-test function from the SciPy package was used for pairwise comparison of the bud count between control and Rhobo6 treated groups.

#### Comparison of Rhobo and Rhobo6 labeling, along with washout

Salivary glands were cultured in Organ Culture Media for 4 days (see above). Two glands from the batch were incubated with Rhobo or Rhobo6 at 5 µM in Organ Culture Media, along with 2 µg/mL Hoechst 33342 (Thermo Fisher Scientific, 62249) to visualize nuclei, and Nucspot650 (Biotium, 41034-T) at 1:500 dilution to visualize dead cells. After 2 h of incubation, glands were mounted as previously described and imaged on a confocal microscope. Subsequently, samples were washed once with fresh warm media and incubated at 37 °C with 5% CO_2_ for 15 minutes, then imaged on a confocal microscope. Two more washes with fresh warm media were conducted during 180 minutes of total incubation time before being imaged a third and final time.

#### Antibody and protein labeling of live salivary glands

Anti-Collagen Type IV antibody (Sigma-Aldrich, AB769) and Anti-Laminin antibody (Sigma-Aldrich, L9393) were fluorescently labeled with Atto647N NHS ester (AAT Bioquest, 2856) following manufacturer instructions. Briefly, antibody vials were adjusted to pH ∼8 via addition of 1:20 v/v of 2 M sodium bicarbonate, pH 9. Atto647N NHS ester was dissolved to a concentration of 10 mM in anhydrous DMSO and incubated with each pH-corrected antibody at a 20:1 molar ratio for 1 h in the dark at room temperature. Free dye was removed using Zeba™ Spin Desalting Columns, 40K MWCO, 0.5 mL (Thermo Fisher Scientific). Note, anti-laminin antibody from the manufacturer contained 1% w/v BSA as a stabilizer, which was fluorophore-labeled alongside the antibody, likely contributing to background signal in live salivary glands. CNA35-GFP was a gift from Jason Northey (UCSF) and was expressed and purified as previously reported^38^. E13 submandibulary salivary glands were isolated as above and cultured for 2 days. Rhobo6 was added at 5 µM alongside ∼10 µg/mL protein label for 1 h. Imaging was performed in the staining solution.

### Mouse excised pancreatic tissue

#### Tissue harvesting

All experiments complied with protocols approved by the Institutional Animal Care and Use Committee (IACUC) at Janelia Research Campus (protocol number 16-142). C57BL/6J mice were obtained from the Jackson Laboratory. All surgical procedures were performed under general anesthesia via administration of ketamine/xylazine (10 mg/kg :10 mg/kg). Krebs-Ringer bicarbonate buffer (KRBH) containing 3 mM D-glucose was injected into the distally clamped bile duct using a 1 mL insulin syringe and a 31G needle. The exsanguinated pancreas was then removed from the peritoneal cavity and cut into 0.5-1.0 cm^3^ pieces in size. Tissue pieces were placed in Krebs-Ringer bicarbonate buffer (KRBH) containing 3 mM D-glucose with 5 µM Rhobo6, 2 µg/mL Hoechst 33342 (Thermo Fisher Scientific, 62249), and/or 10 µg/mL Atto647N-anti-collagen 1 (Novus Biologicals, NB600-408, fluorescently labeled as above). After 1 h of incubation at 37 °C, the tissue was mounted for imaging using double-adhesive imaging spacers and 35 mm glass bottom dishes, as was done with salivary glands (see above).

#### STED microscopy

All experiments complied with protocols approved by the Institutional Animal Care and Use Committee (IACUC) at Janelia Research Campus (protocol number 22-0211). An 8-weeks old C57BI/6J female mouse weighting 20 g was euthanized by cervical dislocation and dissected to extract pancreatic tissue. The tissue sample was cut into ∼5 mm^3^ pieces which were incubated in 3 mL of RPMI-1640 phenol red-free media supplemented with 20 mM HEPES and 5 µM Rhobo6 for 1 h at 37 °C, 95% humidity, 5% CO2. To mount the tissue pieces stably for STED microscopy, samples were placed on a 35mm glass bottom dish (Thermo Fisher Scientific, 150682), and secured by a metal slice anchor (Warner Instruments, 64-1415); 500 µL of Rhobo6-containing media was added on the sample to prevent drying. The mounted sample was imaged within 2 h using a Leica SP8 STED microscope. Both confocal and STED images were acquired of the same field of view to be subsequentially compared. After acquisition, images were denoised (cf. Supplementary Table 1) and an intensity plot was generated in Fiji/ImageJ by extracting pixel value along a line profile averaged over ten pixels.

### Dye administration to mice and live tissue imaging

All experiments complied with protocols approved by the Institutional Animal Care and Use Committee (IACUC) at Janelia Research Campus (protocol number 22-0211). Mice used were between 8- and 12-weeks old C57BI/6J females (Jackson Laboratory) weighting between 20 and 22 grams. 10 µL of Rhobo6 solution at 10 mM in DMSO were diluted with sterile PBS to 100 µL volume to give a 1 mM concentration. A mouse was transferred in an induction chamber and anesthetized with 2.5% isoflurane at a 1.0 L/min oxygen flow rate. The 1 mM Rhobo6 solution was then injected retro-orbitally using a 0.5 mL tuberculin syringe with 27G needle (BD, 305620). Mice were allowed to recover in their cage over 30 mins then euthanized by cervical dislocation. Tissues were dissected onto 35 mm glass bottom dishes (Thermo Fisher Scientific, 150682). For all tissues except for the trachea, the tissues were laid onto the glass whole, for imaging through the fascia into the lumen. The trachea was mounted transversely, with the cross section facing the glass. Notably, mouse urine was bright pink within the 30-minute recovery period prior to euthanasia, indicating that the dye is cleared via glomerular filtration, as expected for a small molecule dye.

#### Second harmonic imaging (SHG) and two-photon excitation autofluorescence microscopy (TPEF)

Jejenum, pancreas and muscle tissues were harvested post Rhobo6 retro-orbital injection as described above. Once tissues were transferred to a 35 mm glass bottom dish (Thermo Fisher Scientific, 150682), they were placed on a two-photon microscope. Once a suitable field of view was identified, images of Rhobo6 fluorescence, TPEF autofluorescence and SHG were acquired (cf. Supplementary Table 1). A control mouse (no Rhobo6 injection), was euthanized and dissected to harvest the same tissue, mounted and imaged in a similar orientation.

### *C. elegans* husbandry and dye administration

Animals were reared at 20 °C on nematode growth media (NGM) plates seeded with HB101 bacteria. Injections were done at room temperature into the syncytium of the distal gonad arm of young to mid-adult N2 or NK2443 *[nid-1(qy38[nid-1::mNG+loxP]) V]* using standard procedures^61^. Animals were injected with approximately 10 pL of PBS or PBS + dye into each gonad arm. After injections, animals were rehydrated in M9 buffer and transferred to NGM plates seeded with HB101 to recover for 30-60 min. Injected animals were anaesthetized with 5 mM sodium azide and mounted on 2% agarose pads and imaged. For Rhobo6 injections, dye aliquots were prepared by diluting with PBS to a final concentration of 100 µM dye and 1% DMSO and stored at −80 °C. Prior to injection, an aliquot was thawed, briefly centrifuged for 5 min at 13000 x g and loaded into the microinjection needle.

### *D. melanogaster* husbandry and dye administration

All flies in this study were raised at 25 °C with a 12-hour light-dark cycle. The following fly stocks were used (stock numbers are from the Bloomington Drosophila Stock Center): Control (w[1118] (#3605)), neuronal cell driver ( w[1118]; P{y[+t7.7] w[+mC]=GMR57C10-GAL4}attP2 (#39171)) and the GFP fluorescent label (w[*]; P{y[+t7.7] w[+mC]= 10XUAS-IVS-Syn21-GFP-p10}attP2^62^. Fly brains were dissected in a chilled modified saline solution of 103 mM NaCl, 3 mM KCl, 5 mM TES, 8 mM trehalose, 10 mM glucose, 26 mm NaHCO_3_, 1 mM NaH_2_PO_4_, 2 mM CaCl_2_ and 4 mM MgCl_2_ at pH 7.4 and placed in a glass bottom 8-well plate (Ibidi, 80807). The fly brains were oriented posterior side down on the plate. Once stuck to the base, the saline solution was removed and Rhobo6 was added at 5 µM. Brains were imaged within 1 h of dissection.

### *D. rerio* husbandry and dye administration

All experiments complied with protocols approved by the Institutional Animal Care and Use Committee (IACUC) at Janelia Research Campus (protocol number 22-0216). Larvae were reared at 28.5 °C in 14-10 light-dark cycles. Zebrafish from 5 d.p.f. were fed rotifers and used for experiments at 8 d.p.f. Zebrafish sex cannot be determined until ∼4 weeks post-fertilization, so experimental animals’ sex was unknown. Fish were embedded in a drop of 2% low-melting-temperature agarose in a glass-bottom petri dish. Agarose surrounding the tail was removed and fish anesthetized with MS-222 (0.16 mg/ml). Water in the sample chamber was then replaced with Rhobo6 solution at 5 µM diluted in fish water containing MS-222. To enable dye delivery an incision in the tail fin was made using a tungsten needle (1 µm tip). Fish were incubated 30 minutes and then imaged.

### *A. thaliana* husbandry and dye administration

*Arabidopsis thaliana* seeds were obtained from the Arabidopsis Biological Resource Center (stock no. CS4004) and placed on a Lloyd & McCown Woody Plant Basal Medium with Vitamins (PhytoTec Labs) agar pad sitting atop a 35 mm glass bottom dish (Thermo Fisher Scientific, 150682). The pad and seeds were incubated at 4 °C for 3 days, then moved to room temperature under lab bench lights for 9 days. Rhobo6 was added at 5 µM to water surrounding the agar pad overnight. Root structures within the agar pad were imaged the following morning.

### Intravital imaging of wild-type and MMTV-PyMT mice

#### Animals and animal care

Animal husbandry of mice was carried out in Laboratory Animal Resource Center (LARC) facilities at UCSF Parnassus in accordance with the guidelines stipulated by the IACUC protocol number AN194983, which adhere to the NIH Guide for the Care and Use of Laboratory Animals. Mice were maintained in pathogen-free, ventilated HEPA-filtered cages under stable housing conditions of 30-70% humidity, a temperature of 20-26°C, and a 12:12 hour dark:light cycle. 10-week-old female FVB/NJ and MMTV-PyMT mice on an FVB/NJ background were used for intravital imaging experiments.

#### Intravital imaging of the mammary gland

Intravital imaging of live animals was conducted according to the IACUC protocol number AN194983 within the Biological Imaging Development Center which was approved by the UCSF IACUC for non-survival experiments. The procedure followed was similar to that described in Dawson *et al.*^63^. Prior to imaging mice were anesthetized in a chamber using oxygen-delivered isoflurane gas (oxygen ∼1 L/min flow rate and isoflurane vaporizer at ∼3%) and Rhobo6 was quickly administered via retroorbital injection before mice were allowed to recover for 15 minutes. Mice were then anesthetized as above before transferring them to a nose cone that was secured to a custom heated microscope stage attachment beneath the microscope objective. Depth of anesthesia was monitored by pedal or toe pinch reflex and breathing rate and adjusted as needed throughout the imaging session. Two small midline incisions were made to open a flap of skin in the mouse which was gently pulled back to reveal the inguinal mammary gland for imaging. A custom metal annulus attached to the stage with height adjustable metal rods was positioned over the mammary gland and pressed to form a seal. A glass coverslip was then affixed to the annulus over the mammary gland with vacuum grease and a drop of water was placed on the coverslip. The stage position was adjusted to place the imaging window directly below the objective lens.

#### Immunofluorescence of wholemounted mammary glands

The same mammary glands that were imaged by intravital microscopy (Figure 5) were resected and fixed in 4% paraformaldehyde for 20 mins. They were then washed with PBS, permeabilized with 0.3% Triton X-100 in PBS for 15 mins before being blocked in PBS with 0.3% Triton X-100, 5% goat serum and 3% bovine serum albumin for 1 h. Mammary glands were then incubated with CNA35-GFP and Phalloidin-647 for 30 mins to stain fibrillar collagens and filamentous actin respectively before being washed with PBS and incubated with DAPI for 10 mins to stain nuclei. A final PBS wash was performed prior to mounting glands with aquamount and a coverslip on a microscope slide. All incubations were done at room temperature and small weights were placed over coverslips to flatten mounted mammary gland tissues as they dried overnight.

## Data availability

Data supporting the findings of this study are available within the article and its Supplementary Information. Unprocessed imaging datasets are available to reviewers on Figshare and will be made publicly available following peer review and publication.

## Extended Data Figures and Figure Captions

**Extended Data Figure 1.**
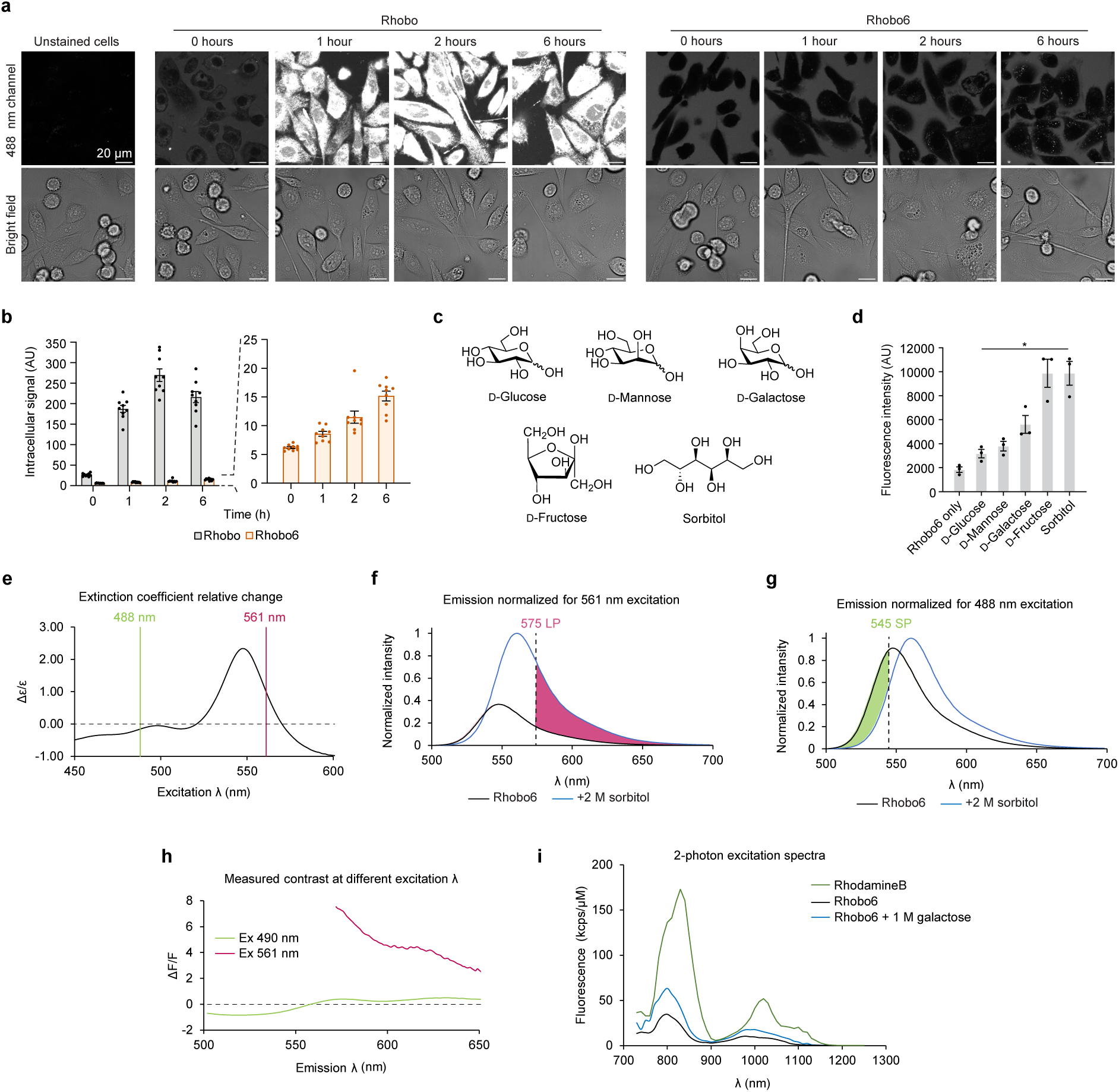
Rhobo6 cell impermeability and additional photophysical characterization. **a,** Comparison between Rhobo and Rhobo6 cell permeability over time. PC3 cells were incubated with Rhobo or Rhobo6 at 5 µM concentration in serum-containing media. Wells were then imaged directly after addition of dye (t = 0 h) and following a 1, 2, and 6 h incubation at 37 °C and 5% CO_2_. Cell surface signal is absent in this experiment due to the presence of serum-containing media (cf. Extended Data Fig. 3d) and excitation at 488 nm (cf. (**e**)-(**h**)). **b,** Quantified intracellular signal for Rhobo and Rhobo6 over time as determined by manually drawn regions of interest of *N* = 9 cells per condition. Error bars represent SEM. **c,** Monosaccharides and monosaccharide analogs used in (**d**). **d,** Quantified fluorescence of Rhobo6 measured by exciting at 555 nm and detecting fluorescence intensity maxima between 570 and 630 nm. All sugars were prepared in PBS solutions at 200 mM with 5 µM Rhobo6, pH 7.3-7.4, and incubated for 1 h at room temperature. *N* = 3, error bars represent SEM. *P* values were determined by unpaired t-test with Welch’s correction, relative to dye only control; \**P* < 0.05. **e**, Relative change in extinction coefficient between bound and unbound Rhobo6 as function of excitation wavelength calculated as (ε_bound_-ε_unbound_)/ε_unbound_. Position of standard laser lines 488 nm and 561 nm are shown to highlight that longer wavelength excitation preferentially excites bound dye, enhancing observed contrast. **f**, Emission spectra of bound and unbound Rhobo6 normalized to 488 nm excitation (cf. *Methods*). A standard emission filter for green fluorescence (545 nm short pass [SP]) will preferentially collect light from unbound dye, reducing observed contrast. **g**, Emission spectra of bound and unbound Rhobo6 normalized for 561 nm excitation. A standard emission filter for red fluorescence (575 nm long pass [LP]) will preferentially collect light from bound dye, enhancing observed contrast. **h**, Measured contrast (calculated as ΔF/F between bound and unbound Rhobo6), as function of emission wavelength for 490 nm and 561 nm excitation. The plot agrees with renormalized spectral data in (**f**)-(**g**), with measured optimal contrast at 561 nm excitation and 570-580 nm emission. **i,** Two-photon excitation spectra for Rhobo6 in 100 mM phosphate buffer, pH 7.4 (unbound state) and 100 mM phosphate buffer containing 1 M galactose, pH 7.4 (bound state). The measured excitation spectrum of Rhodamine B is shown for reference.

**Extended Data Figure 2.**
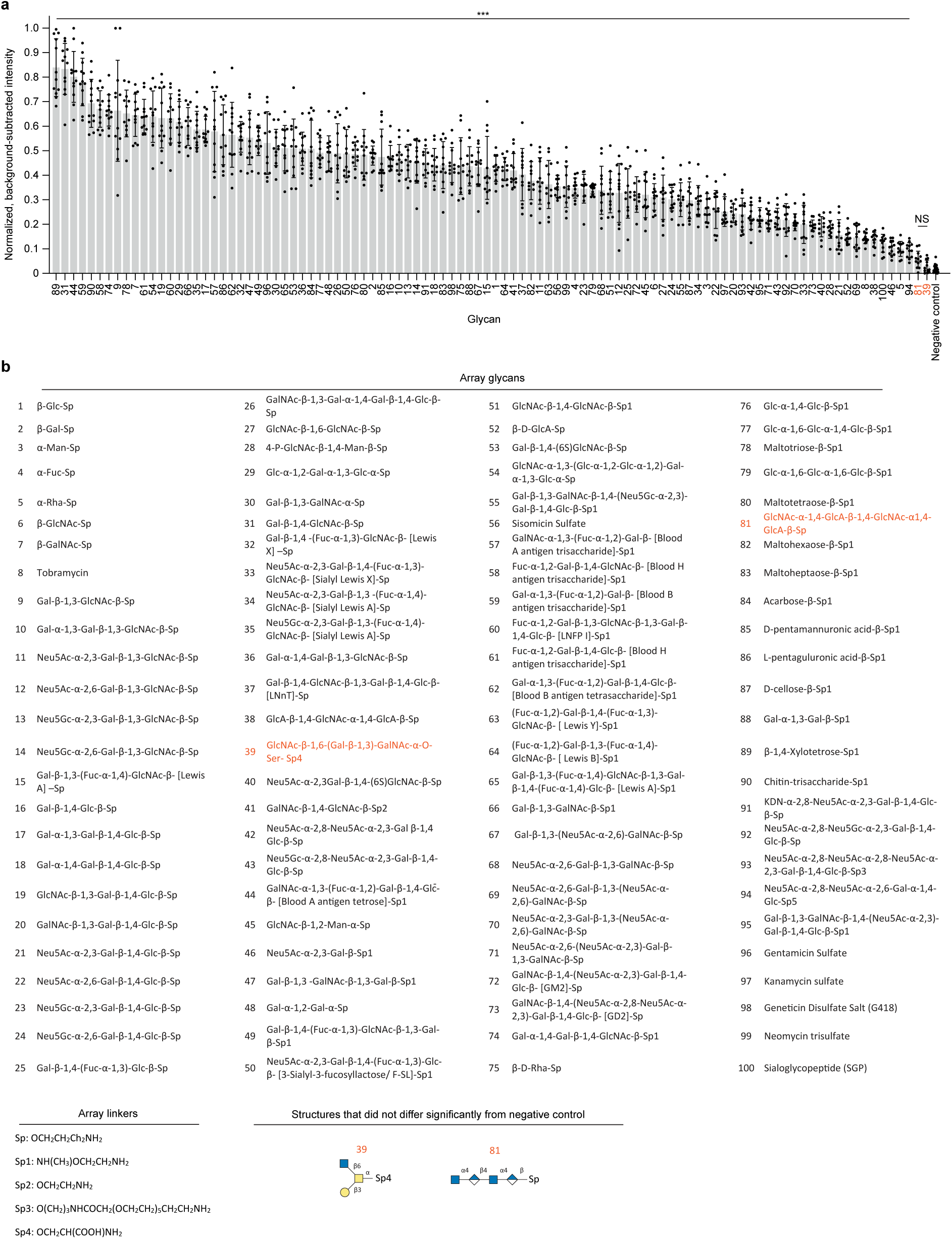
Application of Rhobo6 to a commercial glycan array. **a,** Normalized fluorescence signal of Rhobo6 measured by quantifying three glycan arrays with four glycan replicates each, printed on a single glass slide. Fluorescence signal was background corrected and normalized within each array. *N* = 12 for glycans, *N* = 24 for negative control, error bars represent standard deviation. Statistical significance was determined through Dunnett-corrected t-test for multiple comparisons to a negative control group, with assumption of unequal variance across groups; NS = not significant (orange text), ***P < 0.0005. For all glycans not marked “NS”, average signal was confirmed to be greater than two times the standard deviation of local background. **b,** Array glycans and glycan linkers. Glycans marked “NS” in (**a**) are colored orange. Note that linker “Sp4” carries a carboxylic acid, which is negatively charged at physiological pH.

**Extended Data Figure 3.**
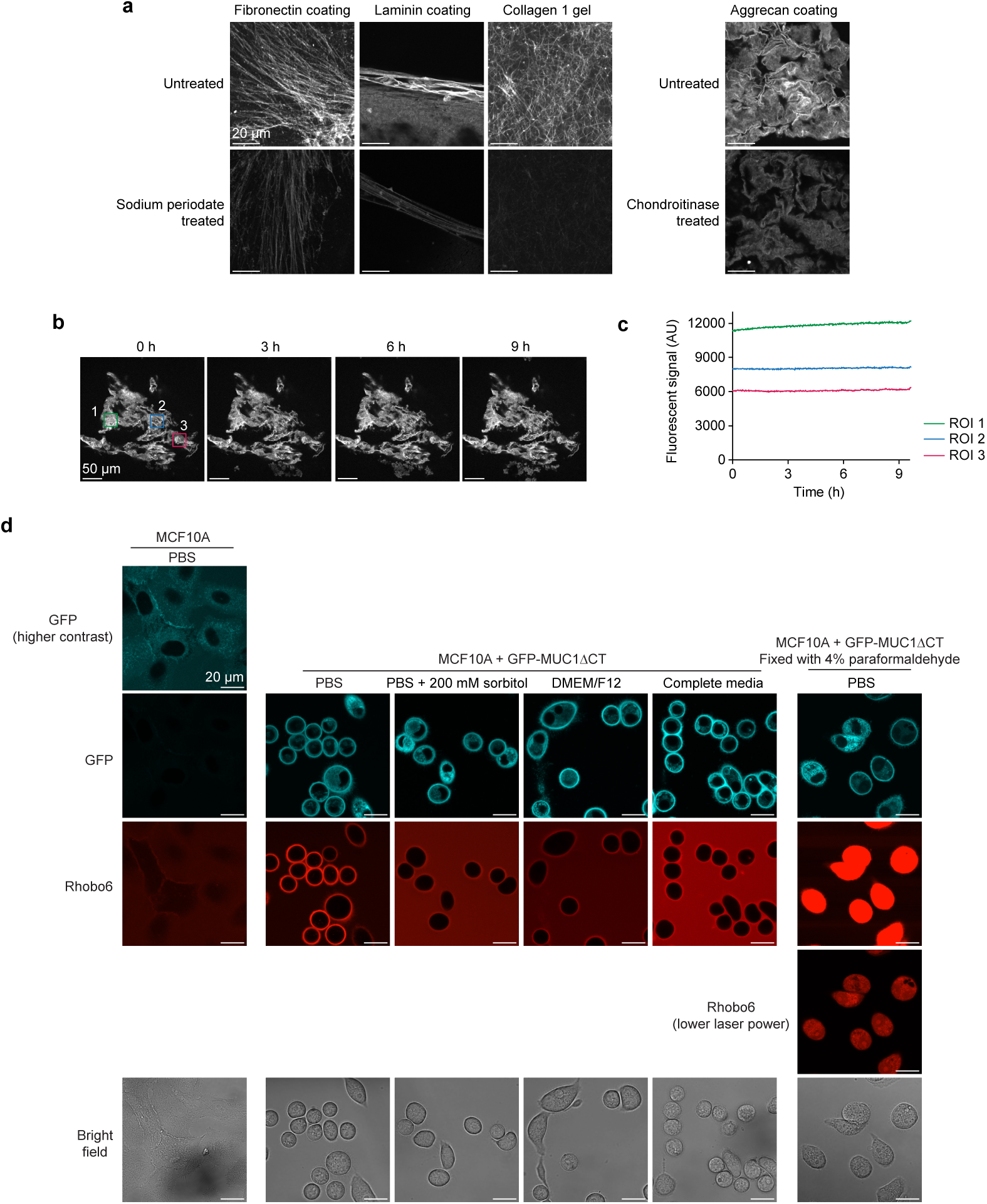
*In vitro* and *in cellulo* Rhobo6 characterization, related to Fig. 2. **a,** Representative field of views from triplicates shown in Fig. 2b. Contrast is normalized within each sample type (columns). Signal intensity is reduced upon treatment, as quantified in Fig. 2c. **b,** Frames extracted from 9.6 h imaging, at one frame per minute, on a coated aggrecan substrate incubated with 5 µM Rhobo6. ROIs analyzed in (**c**) are highlighted in the first frame. **c,** Time trace of signal, calculated as mean intensity within three different manually traced ROIs, from (**b**). Photobleaching was not observed over the course of imaging. **d,** Effect of imaging buffer, mucin expression, and cell fixation on cell surface Rhobo6 signal. MCF10A+GFP-MUC1ΔCT were incubated with doxycycline to induce GFP-tagged mucin expression, and imaged in different buffers containing 5 µM Rhobo6. PBS shows the best signal to background ratio. Cell surface signal can be partially competed upon addition of 200 mM sorbitol (cf. Extended Data Figure 1d), confirming that Rhobo6 binding is diol-dependent and reversible. DMEM/F12 media contains 17.5 mM D-Glucose, likely contributing to a higher background than PBS. Supplementation of DMEM/F12 with the remaining complete media components (cf. *Methods*) results in the lowest observed signal-to-background ratio, likely due to the abundance of glyconjugates in serum. MCF10A cells that were not induced to express GFP-MUC1ΔCT were also imaged in PBS and exhibited dramatically reduced cell surface Rhobo6 labeling, possibly due to the low density of binding sites on these cells. Finally, mucin-expressing cells were imaged after fixation with 4% paraformaldehyde; as a result of the fixation, membrane integrity is compromised, allowing Rhobo6 to accumulate in the cytosol. Rhobo6 is therefore not suitable for use with chemically fixed samples as it will accumulate non-specifically in fixed cells.

**Extended Data Figure 4.**
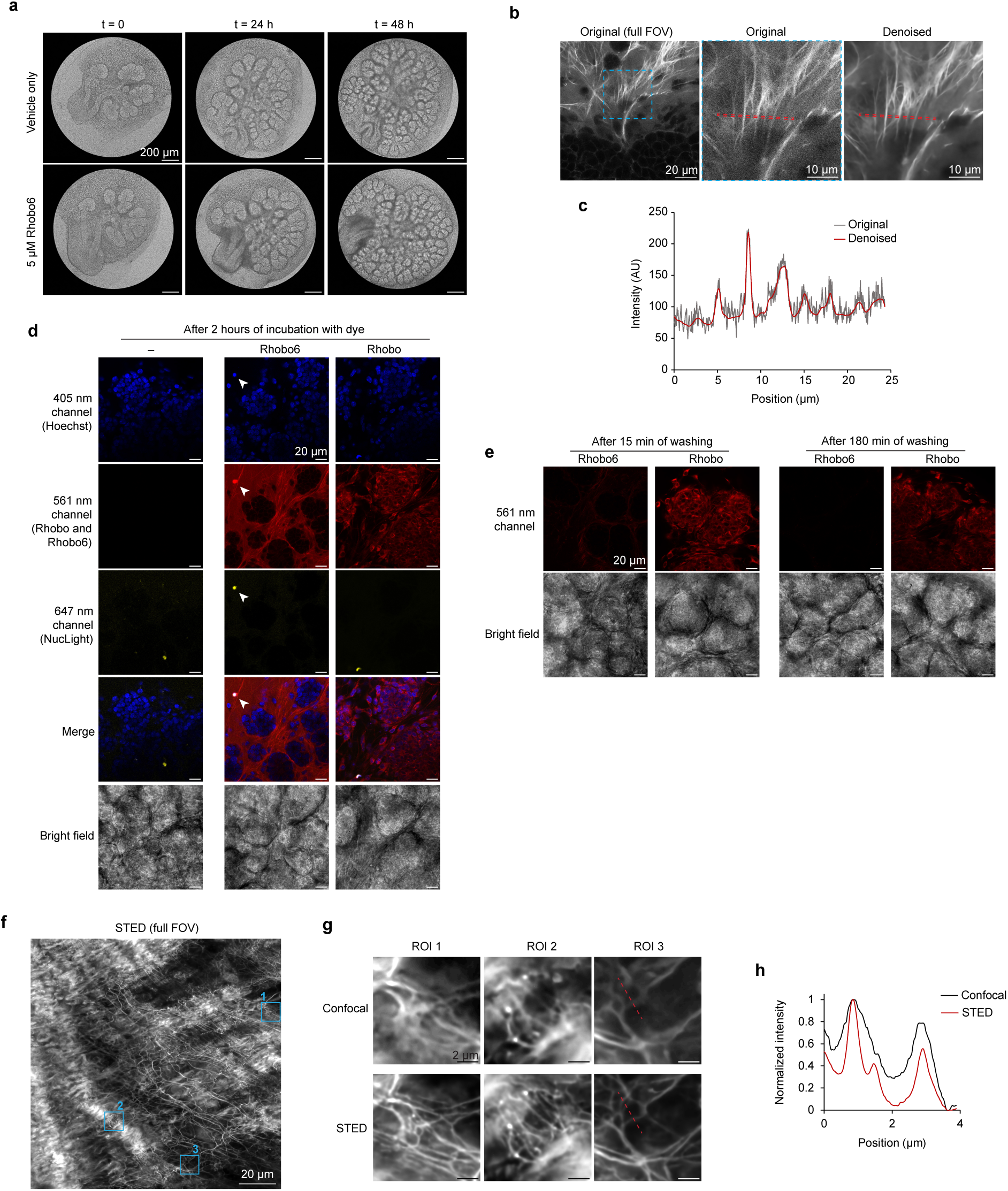
Rhobo6 toxicity, reversibility in comparison to Rhobo, related to Fig. 3. **a,** Brightfield images of a representative pair of glands from Fig. 3b. **b,** Raw image of embryonic salivary gland labeled with Rhobo6 reported in Fig. 3c. Inset shows comparison between raw and denoised dataset, confirming no visual artifacts are introduced in the process. Denoise was performed through Nikon NIS Elements AI Denoise (cf. Supplementary Table 1). Line plot of red region is reported in (**c**). **c,** Line plot of red outlined region from (**b**). Denoising increases signal to noise without altering biological features. **d,** Mouse embryonic salivary glands labeled with 5 µM Rhobo or Rhobo6, along with Hoechst (nuclear stain) and NucSpot650 (dead cell stain). Glands were incubated for 1 h with all probes and imaged. Rhobo6 labeling is confined in the extracellular space, but colocalizes with dead cells due to compromised membrane integrity (arrows). On the other hand, Rhobo labeling accumulates intracellularly in both live and dead cells. **e,** Glands from (**b**) were washed three times over the course of 3 h with DMEM/F12. Rhobo6 labeling was rapidly reversible, while Rhobo labeling was not diminished by washing over the course of the experiment. Rhobo images are contrast normalized across (**d**) and (**e**) and Rhobo6 images are independently normalized across (**d**) and (**e**). Notably, Rhobo is not able to label structures of the extracellular matrix in these glands, likely due to cytosolic sequestration. **f,** Representative STED image of mouse pancreatic ECM obtained by imaging freshly excised tissue labeled with Rhobo6. Depletion was achieved with a 660 nm laser. **g**, Three regions of interest (ROIs) from (**f**), comparing confocal and STED imaging. Contrast is not normalized across imaging conditions and fields of view. Images were denoised (cf. Supplementary Table 1). **h,** Intensity plot of red line from (**g**) displaying the increased resolving power achieved by STED microscopy when compared to diffraction-limited confocal microscopy.

**Extended Data Figure 5.**
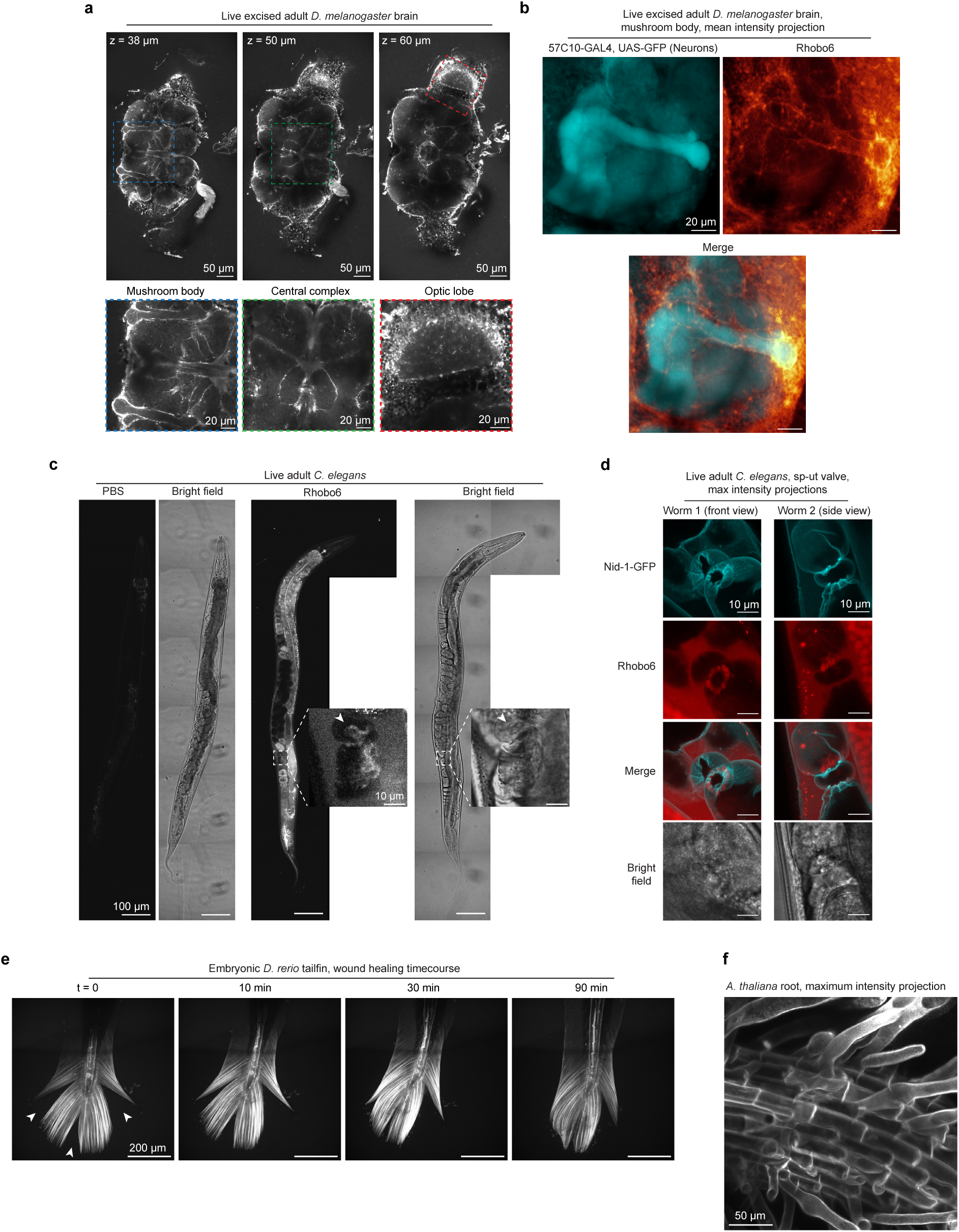
ECM labeling in non-mammalian organisms. **a,** Volume of an adult *Drosophila* brain labeled by bathing with 5 µM Rhobo6 in saline. Imaging at different planes reveals labeling surrounding landmarks in the fly brain, including neuron tracts between the optic lobe and the central brain, the central complex, and the mushroom body. Images were denoised (cf. Supplementary Table 1). **b,** Mean intensity projection of a confocal volume capturing the mushroom body in a *Drosophila* brain. The sample endogenously expressed GFP in neuronal cells and was labeled with Rhobo6 upon dissection (cf. *Methods*). Images were denoised (cf. Supplementary Table 1). **c,** Brightfield and confocal fluorescence images of whole *C. elegans* injected with 10 pL of 100 µM Rhobo6 in PBS containing 1% DMSO. Contrast is normalized between PBS-injected and Rhobo6 injected animals. Inset is a crop and enlargement of the oviduct region. **d,** Max intensity projections of confocal volumes taken at the oviduct of animals endogenously expressing Nidogen-1-mNeonGreen (Nid-1-GFP) to highlight oviduct surfaces, co-localized to Rhobo6 labeling. Rhobo6 signal is enriched at the sp-ut valve within the lumen of the oviduct. Rhobo6 does not label the Nid-1-rich oviduct basement membrane, which is not in contact with the lumen of gonad arm. **e,** Time course of wound healing in zebrafish larvae (8 d.p.f.) incubated with 5 µM Rhobo6 in tank water. Tail nicks (arrows) were necessary for dye delivery. Structures of the tail ECM and notochord are labeled. **f**, Max intensity projection of an *Arabidopsis* root after labeling overnight with 5 µM Rhobo6 in pure water. Root cell surfaces are labeled.

**Extended Data Figure 6.**
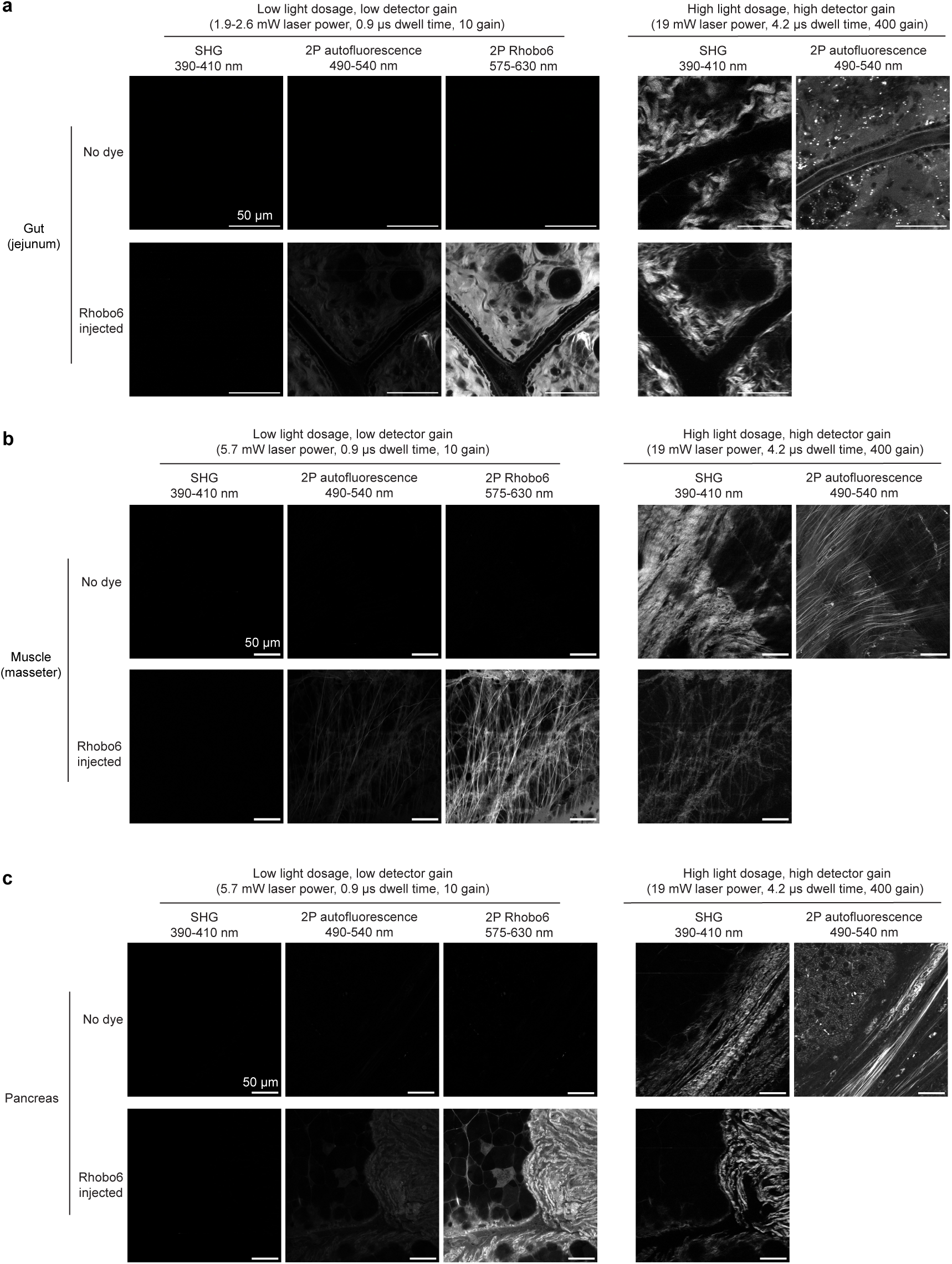
Comparison between 2-photon Rhobo6 imaging and label-free ECM imaging techniques. **a-c**, Gut (jejunum), muscle (masseter), and pancreas from both a control mouse and a mouse retroorbitally injected with 100 nmol of Rhobo6 were imaged using second harmonic generation microscopy (SHG), two photon excitation autofluorescence (2P autofluorescence, also known as TPEF) and two photon excitation fluorescence (2P Rhobo6). Fields of view were imaged with a low light dosage highlighting the efficiency of 2P Rhobo6 compared to SHG and TPEF, followed by a high light dosage for SHG and TPEF, in order to obtain contrast with those methods (cf. Supplementary Table 1). Contrast was normalized across all low dosage images within each tissue type. High dosage SHG images are normalized to each other and high dosage TPEF images are not normalized to any other image. Reported laser power is average measured power at sample plane for 120 fs pulse at 80 MHz repetition rate.

**Extended Data Figure 7.**
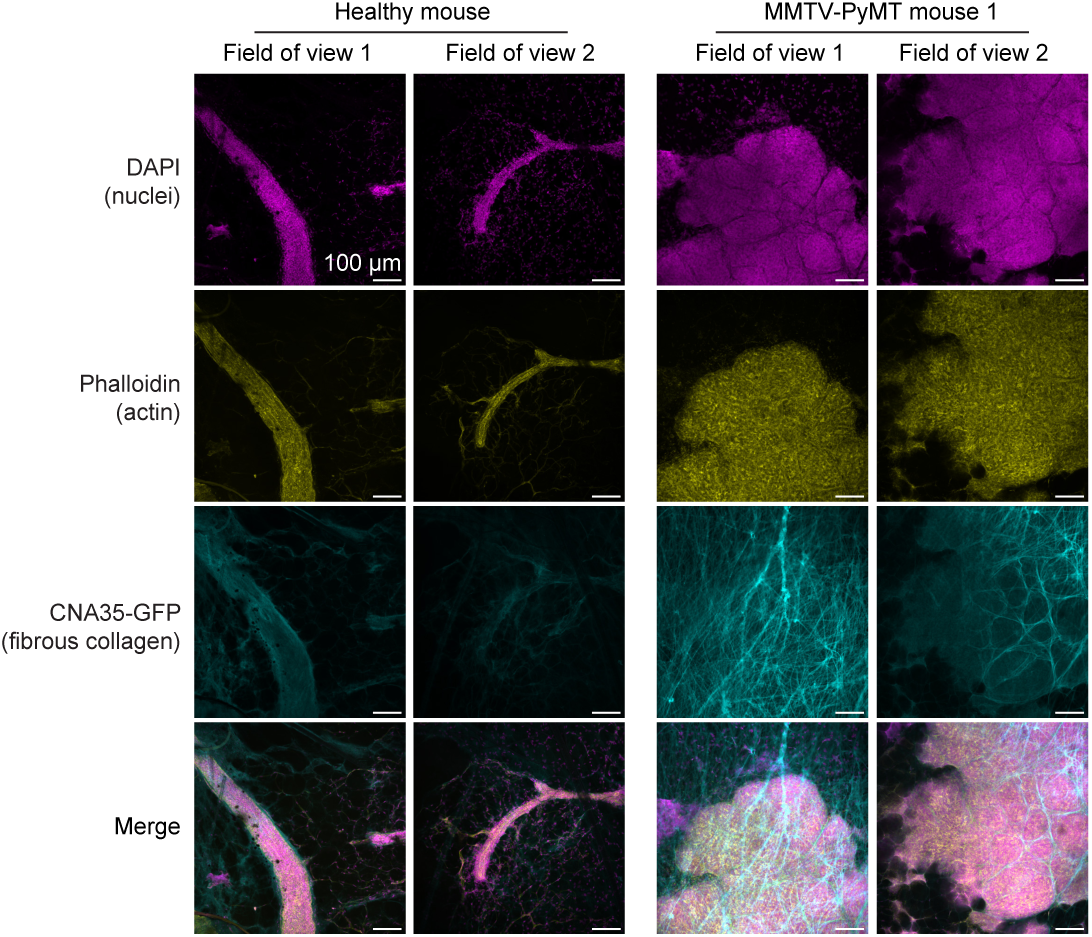
Immunostaining of wild-type and tumor-bearing mouse mammary glands. Immunofluorescence of fixed and wholemounted mammary glands using phalloidin (yellow) and CNA35-GFP (cyan) to mark filamentous actin and fibrillar collagen, respectively, and DAPI to stain cell nuclei (magenta). Two fields of view are shown for each of the mammary glands presented in Fig. 5, resected from the wild-type mouse (left) and MMTV-PyMT mouse 1.

**Extended Data Figure 8.**
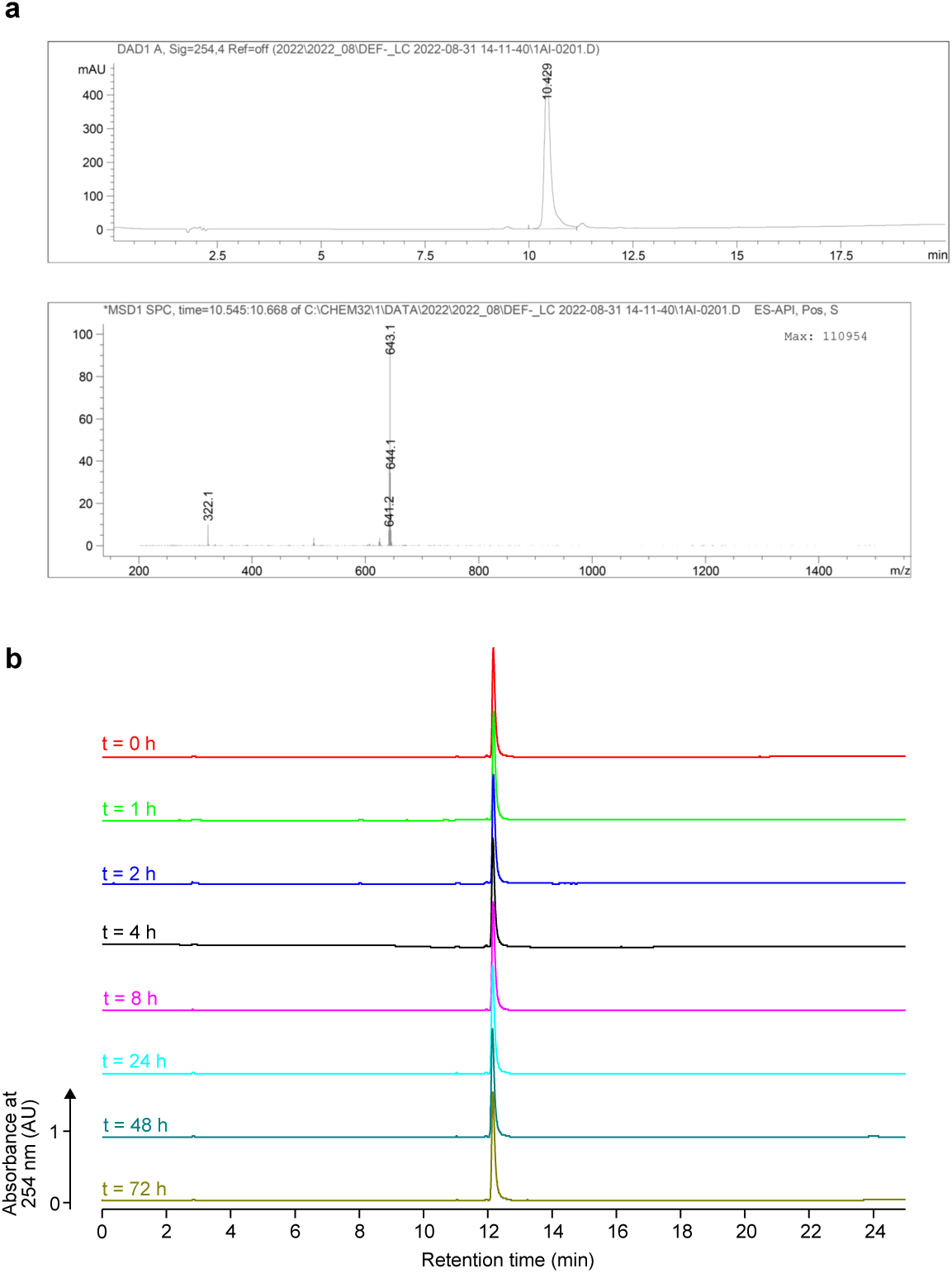
Chemical characterization of Rhobo6. **a,** Analytical liquid chromatography trace with absorbance detection at 254 nm (top) and mass spectrum of main peak (bottom). **b**, Stability of Rhobo6 at room temperature in 1:1 DMSO:PBS over time, as assessed by HPLC with absorbance detection at 254 nm. Calculated Rhobo6 purity ranged from 95-96% over the 72 h period.

